# Human PUM1 and PUM2 exhibit regulation of divergent mRNA targets in male germ cells

**DOI:** 10.1101/760967

**Authors:** Maciej Jerzy Smialek, Erkut Ilaslan, Marcin Piotr Sajek, Aleksandra Swiercz, Damian Mikolaj Janecki, Kamila Kusz-Zamelczyk, Tomasz Wozniak, Maciej Kotecki, Luiza Handschuh, Marek Figlerowicz, Jadwiga Jaruzelska

**Author notes:** authors equally contributed to the manuscript (Erkut Ilaslan, Marcin Piotr Sajek). **To whom correspondence should be addressed:** Jadwiga Jaruzelska, Ph.D., Institute of Human Genetics, Polish Academy of Sciences, Strzeszynska 32, 60-479 Poznan, Poland.

## Abstract

Mammalian Pumilio (PUM) proteins are sequence-specific, RNA-binding proteins with wide-ranging roles, including germ cell development that has functional implications in fertility. Although human PUM1 and PUM2 are closely related to each other and recognize the same RNA binding motif, there is some evidence for functional diversity, particularly related to their roles in fertility. Here, by RNA sequencing (RNA-Seq) approaches, we identified separate mRNA pools regulated by PUM1 and PUM2 proteins in human male germ cells. Using global mass spectrometry-based profiling, we identified distinct PUM1- and PUM2-bound putative protein cofactors, most of them involved in RNA processing. Combinatorial analysis of RNA-Seq and mass spectrometry findings revealed that PUM1 and PUM2 may form distinct RNA-regulatory networks, with different roles in human reproduction and testicular tumorigenesis. Our findings highlight the functional divergence and versatility of PUM paralogue-based post-transcriptional regulation, offering insight into the mechanisms underlying their diverse biological roles and diseases resulting from their dysfunction.

## INTRODUCTION

Posttranscriptional gene regulation (PTGR) is crucial to maintaining cellular proteome homeostasis (Gerstberger, Hafner et al., 2014, Mukherjee, Wessels et al., 2019), disruption of which can cause severe diseases such as cancer and infertility (Fredericks, Cygan et al., 2015). PTGR requires the activity of RNA-binding proteins, such as the widely-studied pumilio (PUM) proteins which are founding members of the PUF (pumilio and fem-3 binding factor) family of eukaryotic RNA-binding proteins. PUM proteins are highly conserved and present in many organisms, from yeast to humans (for review see (Goldstrohm, Hall et al., 2018)). Simultaneous knockout of mouse PUM1 and PUM2 is lethal (Zhang, Chen et al., 2017), indicating their crucial role in development. Posttranscriptional regulation by PUMs is mediated by the conserved C-terminal RNA-binding PUF domain, which is composed of eight tandem repeats (Wang, McLachlan et al., 2002), and binds a specific eight nucleotide sequence 5’-UGUAHAUA-3’, called the PUM-binding element (PBE) that is typically located in the 3’untranslated regions (3’UTR) of target mRNAs. By binding PBEs, PUMs trigger the recruitment of protein cofactors, that together direct selected mRNAs towards post-transcriptional repression or activation (for review see (Goldstrohm et al., 2018)).

Each of the five PUMs in yeast contains a PUF domain that is different in structure from the others, contains between 6-8 tandem repeats and binds to a distinct PBE motif. In this way, each PUM co-ordinately controls the fate of multiple mRNAs sharing a specific PBE motif and which have been found to be functionally related (Gerber, Herschlag et al., 2004). These findings became the basis for the so-called PUM RNA regulon model (Keene, 2007). Considering high structural similarity of PUM1 and PUM2, it is still unresolved whether they form separate regulons in mammals. Although mammalian PUM1 and PUM2 contain nearly identical PUF domains (Spassov & Jurecic, 2003) that recognize the same PBE motif (UGUANAUA) (Galgano, Forrer et al., 2008), there is some evidence for divergent modes of regulation. Examination of interactions between another RNA-binding protein, ARGONAUTE2 (AGO2), and PUM proteins revealed a substantial fraction of nonoverlapping PUM1 and PUM2 mRNA targets (Sternburg, Estep et al., 2018). Therefore, it is possible that PUM1 and PUM2 paralogues are functionally non-redundant and function as distinct RNA regulatory networks (regulons), as previously suggested (Gerber et al., 2004). We recently demonstrated a specific example of functional non-redundancy between PUM1 and PUM2 by showing that while PUM2 induces PBE-dependent repression of the mRNA target *SIAH1*, PUM1 does so in a PBE-independent manner (Sajek, Janecki et al., 2018). Additionally, the regions N-terminal to the PUF domain, which are divergent between PUM1 and PUM2, were reported to contain three unique sub-regions with autonomous repressive activity that may represent an interface for binding protein cofactors since they were not demonstrated to bind RNA (Weidmann & Goldstrohm, 2012).

A number of PUM protein cofactors such as NANOS1, NANOS3 and DAZ family members are associated with male or female infertility in humans (Jaruzelska, Kotecki et al., 2003, Kusz-Zamelczyk, Sajek et al., 2013, Moore, Jaruzelska et al., 2003, Reijo, Lee et al., 1995, Santos, Machado et al., 2014). Also PUM1 itself was found to be important for male and female fertility (Chen, Zheng et al., 2012, Mak, Fang et al., 2016, Xu, Chang et al., 2007). Establishing the mechanisms underlying functional divergence of PUM1 and PUM2 including identification of their protein cofactors may help in understanding their particular roles in human germ cells as well as human infertility, a problem affecting 15% of couples worldwide who are unable to conceive (O’Flynn O’Brien, Varghese et al., 2010). Male infertility in particular impacts 7% of the male population (for review see (Ibtisham, Wu et al., 2017)). Notably, male infertility is a risk factor for developing testis germ cell tumour (TGCT). Testicular cancers are the most frequently diagnosed malignant tumours in young Caucasian males, and their incidence has increased (van de Geijn, Hersmus et al., 2009), highlighting the importance of the human male germ cell context in studying PUM1- and PUM2-controlled regulation. However, the only available germ cell line is TCam-2, which originates from human seminoma, a type of TGCT, and represents male germ cells at an early stage of prenatal development (de Jong, Stoop et al., 2008). The identification of PUM mRNA targets and PUM-interacting proteins had not been previously studied in human germ cells (which would help establishing the mechanisms underlying functional divergence of PUM1 and PUM2 in these cells) and therefore may help in understanding the reasons behind infertility in humans. To the best of our knowledge, the identification of PUM mRNA targets in germ cells has only been studied in the *C. elegans* model (Prasad, Porter et al., 2016). Here, by RNA sequencing and mass spectrometry (MS), distinct mRNA pools and interacting proteins were identified for PUM1 and PUM2 in human germ cells, thereby enabling understanding of PUM functional relevance to fertility.

## RESULTS

### Identification of PUM1 and PUM2 mRNA targets by RIP-Seq

As the first step, to identify human PUM1- and PUM2-bound mRNAs in the TCam-2 cell line, we performed RNA immunoprecipitation (RIP), by combining RNA sequencing (RNA-Seq) and CLIP-Seq protocols. Namely, UV crosslinking step at 254 nm was added to the original RIP-Seq protocol. After verifying the specificity (by performing IP with and without RNase A treatment) of antibodies to PUM1 or PUM2 N-terminal regions and excluding cross-reactions (**Fig. S1A, B** and **Table S1**), IP of these proteins was performed in RNA protecting buffer. Prior to RNA-Seq analysis of PUM-bound mRNAs, we performed RNA-Seq analysis of the TCam-2 transcriptome to use as the reference in RIP-Seq experiments. The RNA-Seq data obtained was in close agreement with the published TCam-2 transcriptome, with a Pearson correlation R^2^ value of 0.957 (**Fig. S2**) (Irie, Weinberger et al., 2015).

The RIP-Seq approach allowed us to identify 1484 and 1133 poly-adenylated RNAs that were significantly enriched (at least two fold) in the anti-PUM1 and anti-PUM2 IPs, respectively, compared to the levels found in IgG IPs and the TCam-2 transcriptome (**Table S2**). Of these, 870 mRNAs were found to specifically bind to PUM1 alone, 519 to PUM2 alone, and 614 (30%) were bound to both PUM1 and PUM2 (**Fig. 1A**) (**Table S2**).

**Fig. 1.**
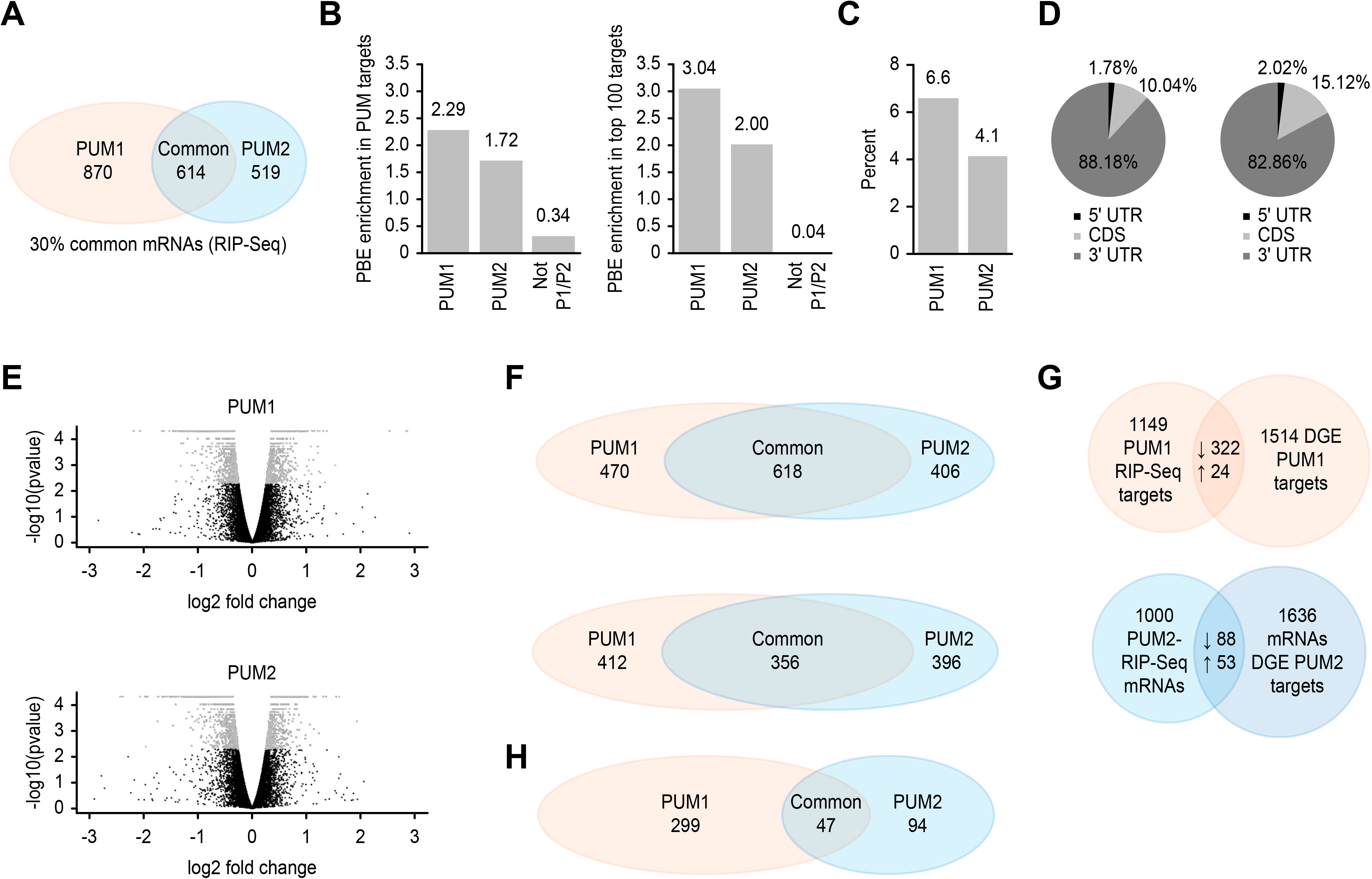
Identification of PUM1- and PUM2 regulated mRNA targets. **A** Venn diagram showing the numbers of PUM1-specific, PUM2-specific and PUM1/PUM2 common mRNA targets from RIP-Seq. **B** PBE enrichment in total PUM1- or PUM2-bound mRNAs (left panel) compared to PBE enrichment in the top 100 mRNAs bound to PUM1 or PUM2 (right panel) calculated by using FIMO software, *P*-value<0.01. **C** Representation of PUM1- or PUM2-bound mRNAs within the whole TCam-2 mRNA transcriptome (%). **D** Diagrams representing PBE motif distribution within the 5’UTR, CDS or 3’UTR of PUM1 (left) or PUM2 (right) bound mRNA targets (FIMO analysis with *P*-value<0.01). **E** Analysis of mRNAs whose expression was significantly changed upon PUM1 or PUM2 siRNA knockdown. Volcano plots representing mRNAs under PUM1 or PUM2 activation (left side of each plot) or repression (right side of each plot). Grey dots on the top of each plot represent changes in mRNA level with *P*-value <0.05 (considered statistically significant). **F**. Venn diagram representing the numbers of mRNAs repressed (upper graph) or activated/stabilized (lower graph) by PUM1, PUM2 or both (grey dots from **Fig. 1E**). The Venn diagram represents the number of mRNAs increased (upper graph) or decreased (lower graph) upon siRNA knockdown of PUM1 (pink), PUM2 (blue) or both. **G** Venn diagrams showing mRNAs that were regulated by PUM proteins (as identified by both the RIP-Seq approach and siRNA-Seq approach); PUM1-regulated (upper panel), PUM2-regulated (middle panel). ↑ activated, ↓ repressed mRNAs **H**. Venn diagram representing the numbers of mRNAs regulated by PUM1, PUM2 or commonly regulated based on data presented in G.

Although it was previously established that each human PUM paralogue specifically recognizes the PBE motif UGUANAUA (Galgano et al., 2008), the PBE motif UGUAHAUW (H stands for A, C or U while W for A or U) was found in a recent study to be more accurate for PUM1 and PUM2 (Bohn, Van Etten et al., 2018). Therefore, in this study, mRNAs identified by RIP-Seq were screened for the presence of the UGUAHAUW motif. We found that on average, PUM1-bound mRNAs contain 2.29 UGUAHAUW motifs/sequences, while PUM2-bound mRNAs contain 1.72 (**Fig. 1B** left panel). The same analysis when performed for the 100 most enriched PUM1 and PUM2 targets, revealed that the motif frequency was higher, with PBE content of 3.04 and 2.00 for PUM1 and PUM2, respectively (**Fig. 1B** right panel). In contrast, in non-specifically bound mRNAs (those present in immunoprecipitates but that were not significantly enriched (<2x) in comparison to non-immune serum and the TCam-2 transcriptome), PBE motif occurrence was significantly lower (0.34/sequence for all RIP-Seq identified and 0.04/sequence for top 100 targets with a higher occurrence in anti-IgG than in anti-PUM immunoprecipitates) (**Fig. 1B**). We observed that altogether ~6.6% of the TCam-2 transcriptome presented PUM1-bound mRNAs, and ~4.1% presented PUM2-bound mRNAs and almost all of them contained PBEs (**Fig. 1C**). We next checked for PBE motif localization within each PUM1 and PUM2 target mRNA to determine the percentage of target mRNAs that harboured PBEs in the 3’UTR or in other locations. We found that PBEs were mostly located in the 3’UTR (88% and 83% for PUM1 and PUM2, respectively), less frequently in CDS (10% and 15% for PUM1 and PUM2, respectively) and rarely in the 5’UTR (1.8% and 2.0% for PUM1 and PUM2, respectively) (**Fig. 1D**). These numbers are in agreement with those obtained from HeLa cells (Galgano et al., 2008) as well as recent data from HEK293T cells (Bohn et al., 2018).

To address whether PUM1- and PUM2-bound mRNAs selected by the RIP-Seq approach represented similar or different cellular functions, we performed Gene Ontology analysis (**Fig. S3, Table S3**). Since we obtained similar numbers of mRNA bound to PUM1 and PUM2 (1484 and 1133, respectively), we were able to use GO BiNGO plug-in in Cytoscape platform (Maere, Heymans et al., 2005) to compare the biological processes and molecular functions of these two groups in an unbiased manner using our TCam-2 transcriptome as a background. We found that while the majority of targets represented overlapping biological processes (BP) and molecular functions (MF), some were related only to PUM1 targets, e.g., chromosome organization, positive regulation of transcription, chromatin modification (BP from **Fig. S3A**), transcription regulator activity, GTPase activator activity and protein binding (MF from **Fig. S3B**). Some other functions were related only to PUM2 targets, e.g., cell cycle, organelle organization, M phase (BP from **Fig. S3A**), motor activity, helicase activity and cytoskeletal protein binding (MF from **Fig. S3B**).

### Differential gene expression analysis upon PUM1 or PUM2 siRNA knockdown

Since mRNA binding alone does not imply regulation by RBPs, and PUM1-repressed mRNAs have been reported to undergo degradation (Morris, Mukherjee et al., 2008) or activation (Bohn et al., 2018), we next sought to identify those mRNAs whose expression was modified upon *PUM1* or *PUM2* gene knockdown. At 72 h after TCam-2 cell transfection with *PUM1* or *PUM2* siRNA, when silencing efficiency was the highest for both paralogues (**Fig. S1C, D**), RNA was isolated and RNA-Seq analysis was conducted, we refer to this approach as siRNA-Seq from hereon. This analysis revealed 1088 genes with higher expression and 768 genes with lower expression upon PUM1 knockdown, 1024 genes with higher expression and 752 genes with lower expression upon *PUM2* gene knockdown (with the adjusted *P*-value<0.05) (**Fig. 1E, F, Table S4**). RNA-Seq analysis revealed that among these, 470 genes were specifically repressed and 412 genes were specifically activated by PUM1 alone, 406 genes were specifically repressed and 396 genes were specifically activated by PUM2 alone, 618 genes were repressed by both PUM1 and PUM2, and 356 genes were activated by both PUM1 and PUM2 (**Fig. 1E, F, Table S4**). Gene Ontology analysis of mRNAs up- and downregulated upon PUM1 and/or PUM2 siRNA knockdown (PUM1 or PUM2 siRNA-Seq) is presented in **Fig. S4**, and shows that most of biological processes and molecular functions of siRNA-Seq PUM1 and PUM2 targets overlap.

### Selection of PUM1- and PUM2-regulated mRNA targets based on RIP-Seq and siRNA-Seq analysis

For the identification of PUM1- and PUM2-regulated mRNA targets, we used the following two selection criteria: #1 binding to PUM1 or PUM2 as detected by RIP-Seq (genes listed in **Table S2**) and #2 down- or upregulation of mRNA levels upon *PUM1* or *PUM2* siRNA knockdown and RNA-Seq (siRNA-Seq, genes listed in **Table S4**). The simultaneous use of both criteria provided us with 346 (322 repressed and 24 activated) PUM1-regulated (**Fig. 1G** upper panel, **Table S5**), 141 (88 repressed and 53 activated) PUM2-regulated (**Fig. 1G** lower panel, **Table S5**) mRNAs. Additionally, by using these two criteria, we found that the number of mRNAs shared by PUM1 and PUM2 was reduced to 10% (47 common mRNAs) (**Fig. 1H**) compared to the 30% seen by RIP-based selection (**Fig. 1A**).

The mRNAs regulated by both PUM1 and PUM2 represented only 1.35% and 0.62% of the TCam-2 transcriptome, respectively (**Fig. 2A**), compared to 6.56% and 4.14% PUM1- or PUM2-bound mRNAs (**Fig. 1C**). To further validate the mRNA pools that we considered to be regulated by PUM1 and PUM2 (**Fig. 1G**), we analysed their PBE-motif content. We found that the number of mRNAs containing at least one PBE reached nearly 100% (96.82% for PUM1 and 99.76% for PUM2) (**Fig. 2B**). This is significantly higher than the PBE content in RIP-Seq- or siRNA-Seq-identified mRNAs (RIP-Seq PUM1 90.77%; RIP-Seq PUM2 85.94%; siRNA-Seq PUM1 59.68%; siRNA-Seq PUM2 57.50%). This result additionally validated our approach. PBE motif distribution in regions of siRNA-Seq PUM mRNA targets is shown in **Fig. 2C**. We also found that in the case of mRNAs under positive regulation by PUM1 and/or PUM2, PBE motif frequency in the 5’UTR was almost 4 times higher than in mRNAs pools under PUM1 and/or PUM2 repression (**Fig. 2D**), suggesting that the 5’UTR sequence is more frequently used in the case of mRNAs activated/stabilized by PUMs than in repressed mRNAs.

**Fig. 2.**
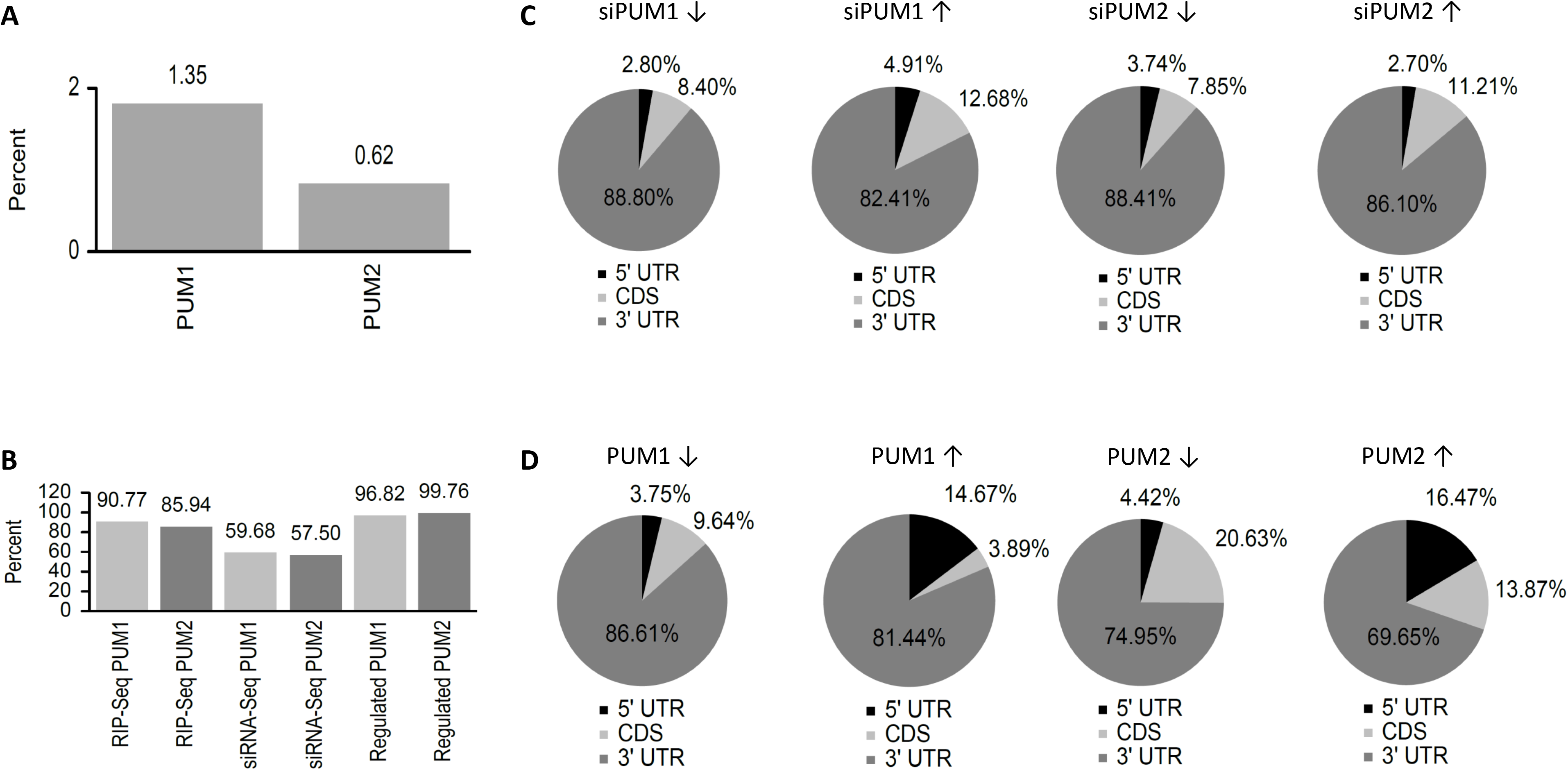
Content of PUM-regulated mRNAs within the TCam-2 cell transcriptome and comparison of PBE content and distribution in PUM-regulated mRNA pools identified by RIP-Seq, siRNA-Seq, and by combined RIP-Seq/siRNA-Seq. **A** Content of PUM1- and PUM2-regulated mRNAs in the TCam-2 mRNA transcriptome (%). **B**. Content of mRNAs containing at least one PBE at three steps of selection: 1/ RIP-Seq approach, 2/ PUM1 or PUM2 siRNA knockdown and 3/ combined RIP-Seq/siRNA-Seq (regulated). **C** PBE distribution in particular regions of mRNAs whose level was changed upon PUM1 (repressed - first or activated - second from the left) and PUM2 (repressed - third or activated - fourth from the left) siRNA knockdown, respectively. **D** PBE distribution in particular regions of regulated mRNAs by PUM1 (repressed - first or activated - second from the left) and PUM2 (repressed - third or activated - fourth from the left), respectively.

Gene ontology analysis revealed that most of the biological processes and molecular functions of mRNAs regulated by PUM1 and PUM2 are different. While PUM1 regulated targets are involved e.g., in mitotic cell cycle checkpoint, regulation of transcription, developmental process, chromosome localization (BP from **Fig. S5A**), small GTPase regulator activity and protein binding (MF from **Fig. S5B**), PUM2 regulated targets are involved e.g., in negative regulation of cell division (BP from **Fig. S5A**), signal transducer activity, MAP kinase activity, kinase activity and transferase activity (MF from **Fig. S5B**). GO analysis revealed also a minority of molecular functions involving both, PUM1 and PUM2 regulated mRNAs, e.g., nucleoside-triphosphatase regulator activity, enzyme regulator activity and nucleoside binding (MF from **Fig. S5B**).

### Global profiling reveals many putative protein cofactors bound by PUM1 and PUM2 to be RNA-binding proteins

Our result showing that PUM1 and PUM2 share only ~10% of their mRNA targets (**Fig. 1H**) was surprising given that PUM1 and PUM2 recognize the same UGUAHAUW motif (Bohn et al., 2018) and show remarkably high similarity in binding potential across 12,285 sequences, as shown by quantitative analysis of RNA on a massively parallel array (RNA-MaP) (Jarmoskaite, Denny et al., 2019). Since N-terminal regions are known to be structurally divergent (Spassov & Jurecic, 2003), they might function differently in PUM1 and PUM2, for example, by binding different sets of protein cofactors. We hypothesized that PUM1 and PUM2 discriminate between specific mRNA targets *in vivo* by interacting with unique protein cofactors. To test this hypothesis, we performed anti-PUM1 and anti-PUM2 co-IP experiments followed by mass spectrometry (MS) to identify PUM1- and PUM2-binding proteins in TCam-2 cells. We identified 27 PUM1-, 13 PUM2- and 7 PUM1/PUM2-interacting proteins, all of which required the presence of RNA for binding (**Fig. 3A**). They all represent known RNA-binding proteins (RBPs), which supports our results. However, we also identified 15 PUM1-, 34 PUM2-, and 15 PUM1/PUM2-interacting proteins that interacted in an RNA-independent manner (**Fig. 3B**). Twenty-eight of these PUM-interacting proteins were identified both in the presence and absence of RNA (**Table S6** and **S1**). Taken together, we identified 27 PUM1-, 30 PUM2- and 25 PUM1/PUM2-bound protein interactors (**Table S6** and **S1**). Interestingly, 54 among 82 PUM1- and PUM2-bound proteins identified in this study are known RBPs, and for 26 of them, a specific RNA-binding motif has already been established using Photoactivatable Ribonucleoside-Enhanced Crosslinking and Immunoprecipitation (PAR-CLIP) or RNA competing methods (Hafner, Landthaler et al., 2010, Ray, Kazan et al., 2013) (**Table S6** and **S1**). Proteins bound to PUM1 were mostly different from those bound to PUM2, with the majority of them being functionally involved in RNA binding, regulation and processing.

**Fig. 3.**
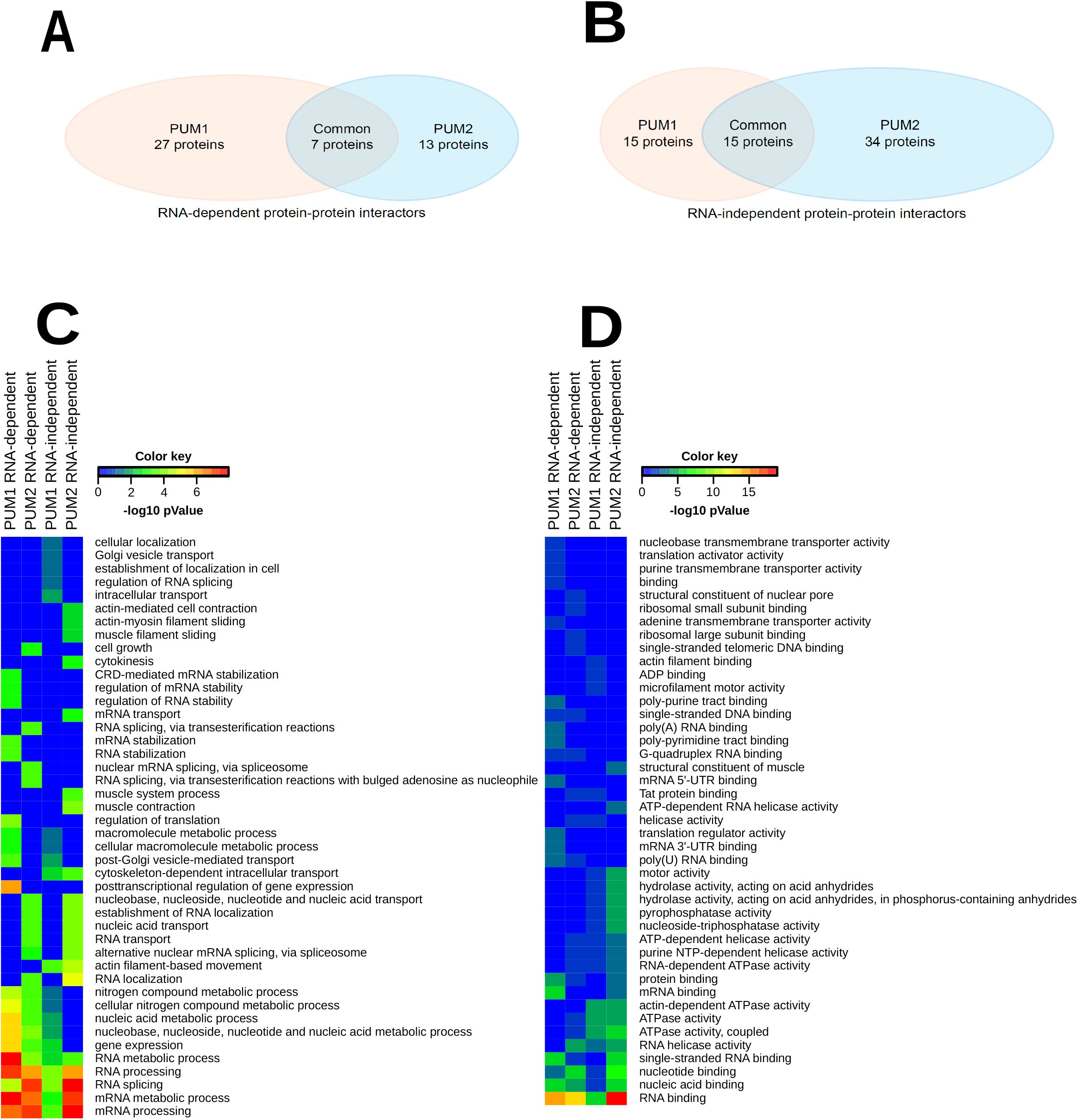
Identification of proteins interacting with PUM1, PUM2 or both, identified by co-immunoprecipitation (co-IP) and mass spectrometry (MS). **A** Proteins interacting with PUM1, PUM2 or both via RNA (co-IP without RNase A treatment). **B** Proteins interacting with PUM1, PUM2 or both, independently of RNA (co-IP with RNase A treatment). **C** Heatmap of BiNGO analysis of TOP20 Biological Processes of PUM1 or PUM2 bound proteins in RNA dependent or independent manner. **D** Heatmap of BiNGO analysis of TOP20 Molecular Functions of PUM1 or PUM2 bound proteins in RNA dependent or independent manner. The detailed results of the GO analysis are presented in **Table S3**.

### Binding motifs of PUM1- and PUM2-bound RBPs are enriched in PUM-regulated mRNAs

To check whether PUM1- and PUM2-bound RBPs could potentially cooperate with PUM in the selection of specific mRNA targets for regulation, we first checked whether the binding motifs corresponding to these RBPs co-occur with PBEs in PUM1- and PUM2-regulated mRNA targets. To this end, we performed an analysis of binding motif enrichment for 10 PUM1-specific (**Fig. 4A**) and 8 PUM2-specific RBPs (**Fig. 4B**) in PUM1 or PUM2-regulated mRNA targets, respectively. We found that RNA binding motifs (**Fig. 4D**) for 6 out of 10 PUM1 bound RBPs – IGF2BP3, YBX1, NUDT21, IGF2BP1, PABPC4 and CPSF7, but not FUS, LIN28A, HNRNPK and CPSF6, were highly enriched in PUM1 regulated mRNAs (**Fig. 4A**). In the case of PUM2, we found that RNA binding motifs (**Fig. 4D**) for 7 out of 8 PUM2 bound RBPs – PTBP1, G3BP2, G3BP1, HNRNPF, FMR1, SRSF7 and SRSF1 but not HNRNPA2B1, were highly enriched in PUM2 regulated mRNAs (**Fig. 4B**). Additionally, we also performed analysis of motif enrichment for 7 common PUM1- and PUM2-interacting RBPs in mRNAs regulated by both PUM1 and PUM2 (**Fig. 4C**) and found that RNA binding motifs (**Fig. 4D**) for all common PUM1 and PUM2 bound RBPs – SFPQ, FXR1, FXR2, NCL, HNRNPA1, MATR3 and PABPC1, were highly enriched in both, PUM1 and PUM2 regulated mRNAs. Motif enrichment was evaluated relative to motif enrichment in nonregulated mRNAs set as 1 (**Fig. 4A-C** dashed lines). The high enrichment of RBP motifs that we found is an additional indication for these RBPs to be putative PUM1 or PUM2 protein cofactors in the regulation of their mRNA targets.

**Fig. 4.**
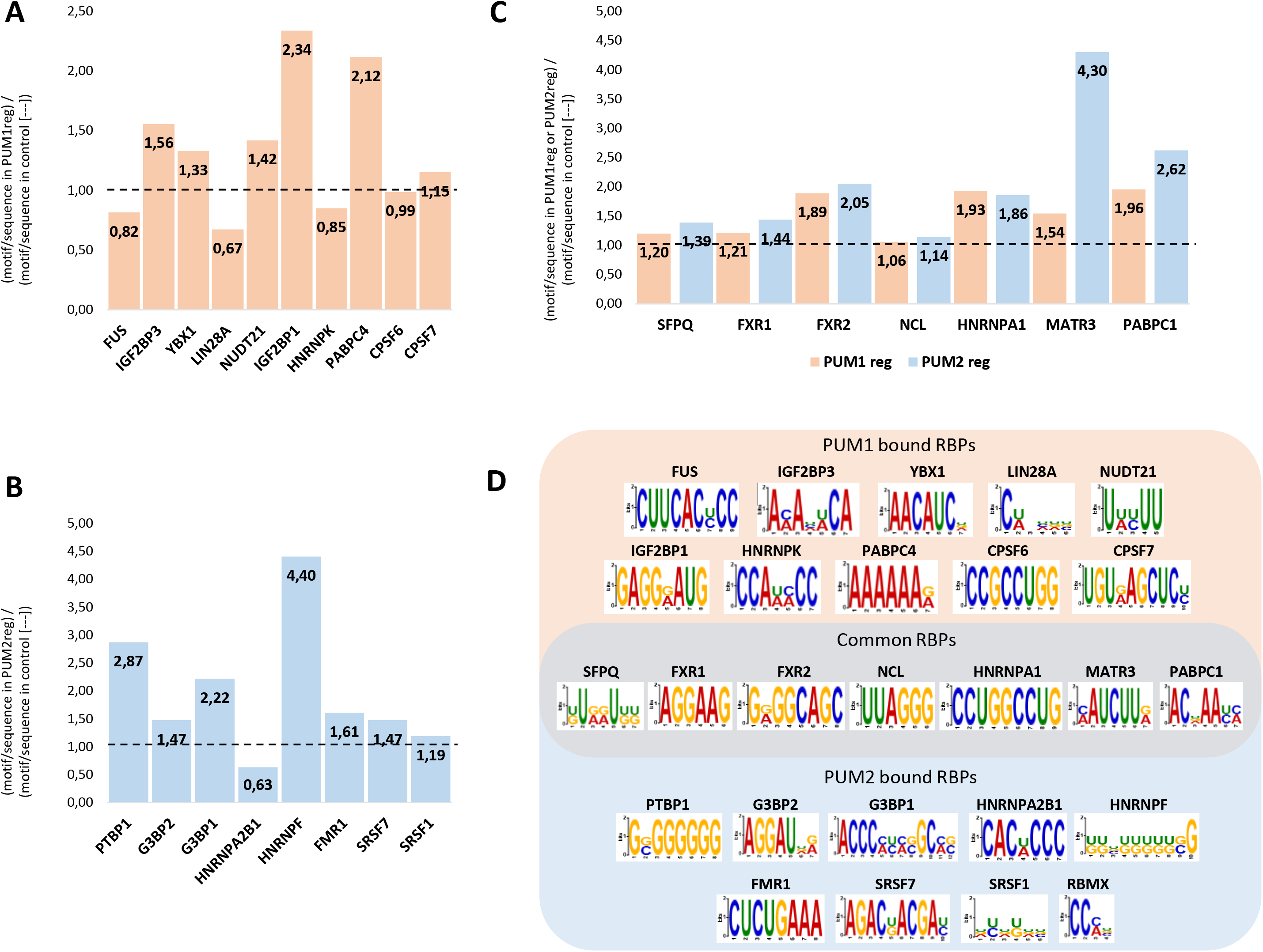
Enrichment of binding motifs for PUM1- or PUM2-interacting RBPs identified in each mRNA regulated by PUM1 or PUM2 in TCam-2 cells. The entire sequence of each mRNA was searched for these motifs. Each enrichment analysis was performed using FIMO software with a threshold set at *P*-value<0.01. Motif enrichment values of particular RBPs in mRNA from the TCam-2 transcriptome not regulated by PUMs were set at 1, and this was the baseline control (shown with dashed line). **A** Histograms representing the enrichment of binding motifs of PUM-interacting RBPs within mRNAs regulated by PUM1 are in pink. **B** Histograms representing enrichment of binding motifs of PUM2-interacting RBPs in mRNAs regulated by PUM2 are in blue. **C** Enrichment of binding motifs of RBPs interacting with both PUM1 and PUM2 in mRNAs regulated by both PUM1 (pink) and PUM2 (blue). **D** Motifs corresponding to RBPs (and used in this analysis) interacting with PUM1 and/or PUM2 were generated using FIMO software.

### PUM1 and PUM2 Form separate regulons in TCam-2 cells

To further explore potential functional specificities between PUM1 and PUM2, we combined the above RIP-Seq (**Fig. 1**), siRNA-Seq (**Fig. 1**), co-IP/MS (**Fig. 3**), RNA binding motif enrichment (**Fig. 4**) and GO analysis data. This combined analysis was based on the assumption that an mRNA containing a binding motif for a specific PUM protein cofactor, the frequency of which is significantly higher (above average) than in the control mRNA dataset, is co-regulated by that protein cofactor (**Fig. 3** and **4**) (**Table S7**). The main findings are as follows: First, there are separate PUM1 and PUM2 regulatory networks (regulons). Second, PUM1 and PUM2 may cooperate with varied components to regulate different pathways. As examples, PUM1 and IGF2BP1 may co-regulate mRNA sub-pools involved in intracellular lipid transport; PUM1 together with PABPC4 and MATR3 may co-regulate mRNAs involved in epidermal growth factor receptor signalling pathway; and PUM1 together with PABPC1 and PABPC4 may co-regulated mRNAs involved in negative regulation of binding. On the other hand, PUM2 and SRSF1 may co-regulate endothelial cell development; PUM2 together with SFPQ and SRSF7 – establishment of cell polarity and cell morphogenesis involved in neuron differentiation; PUM2 together with G3BP2, HNRNPA1, FXR2 and NCL may co-regulate mRNAs involved in regulation of Rho protein signal transduction. PUM2 and MATR3 may co-regulate mRNAs involved negative regulation of cell development; PUM1 together with IGF2BP3 may co-regulate mRNAs involved in regulation of cell division; PUM1 together with PABPC4, IGF2BP3, YBX1, NUDT21 and MATR3 may co-regulate mRNAs involved in histone lysine methylation. Third, although PUM1 and PUM2 form separate regulons, they may cooperate in the regulation of some common mRNA targets, which are involved in the same biological processes (**Fig. 5**).

**Fig. 5.**
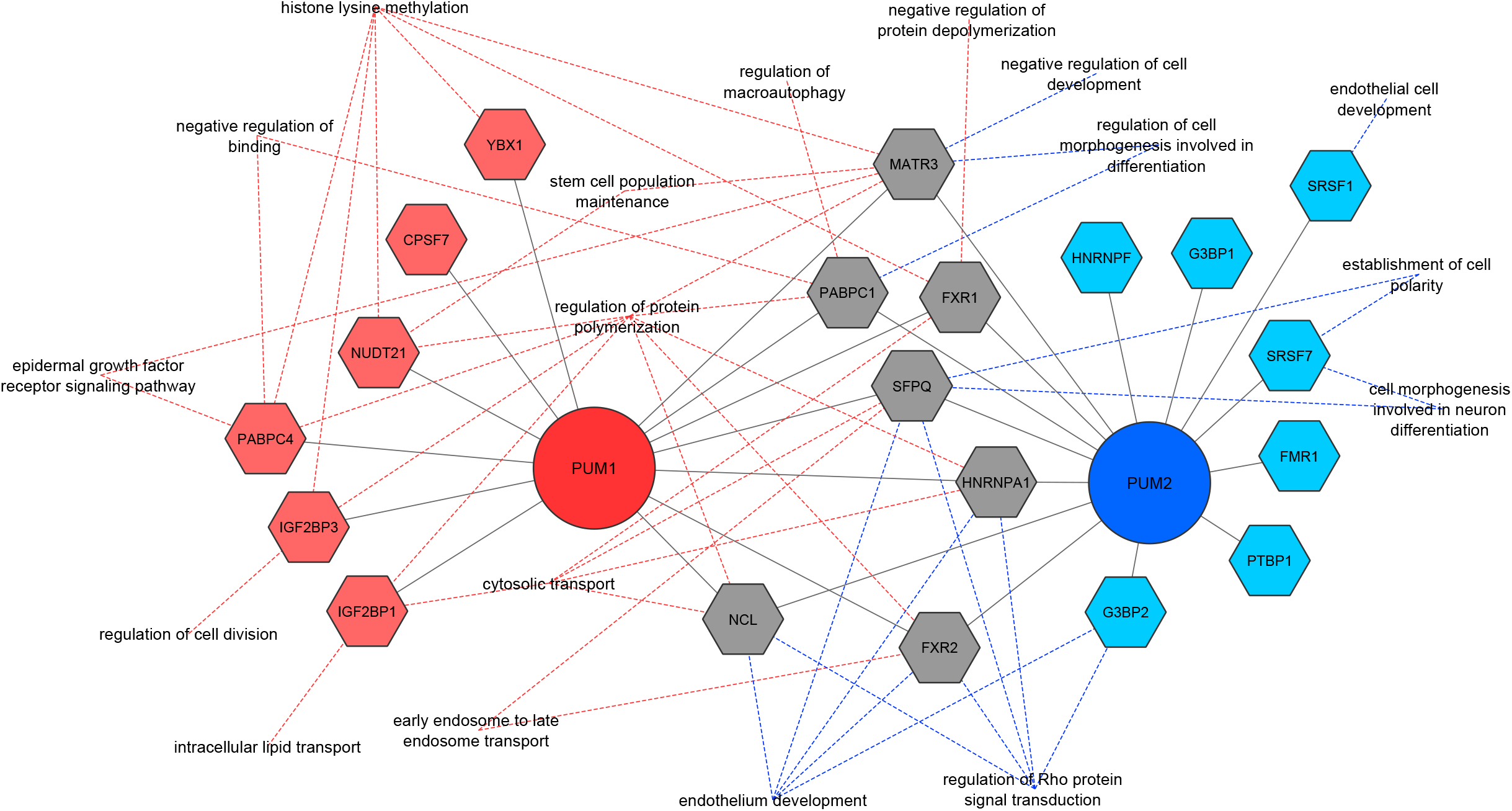
Model of PUM1 and PUM2 regulatory units (regulons) in TCam-2 cells built using the Cytoscape platform. PUM1and PUM2 are represented by red and blue circles, respectively, while PUM1, PUM2 and PUM1/PUM2 interacting RBPs by red, blue or grey diamonds, respectively. Continuous lines represent interactions of PUM1 and PUM2 with RBP putative cofactors. Dashed lines represent functions (biological processes from BiNGO analysis) of groups of mRNAs regulated by PUM1 (dashed red lines) and PUM2 (dashed blue lines) with high enrichment (above average) of binding motifs for particular RBPs in mRNAs regulated by PUM1 or PUM2.

### Several mRNAs highly expressed in TCam-2 cells compared to somatic gonadal tissue are regulated by PUM proteins

Of the many cellular processes regulated by PUM proteins, those involved in germ cell development are of particular interest due to implications to understand infertility in humans (for review see (Goldstrohm et al., 2018)). Therefore, we determined which genes highly and selectively expressed in germ cells are under PUM1 and/or PUM2 regulation. To this end, we first identified genes whose expression in germ cells was at least six times higher than in human testis somatic gonadal tissue by comparing TCam-2 transcriptomic data with the previously published transcriptome of human testis somatic gonadal tissue (Irie et al., 2015). This comparison identified 565 genes highly expressed in TCam-2 (**Fig. 6A**), including 22 regulated by PUM proteins. Specifically, we identified 13 genes regulated by PUM1 alone, 5 genes regulated by PUM2 alone and 4 by both.

**Fig. 6.**
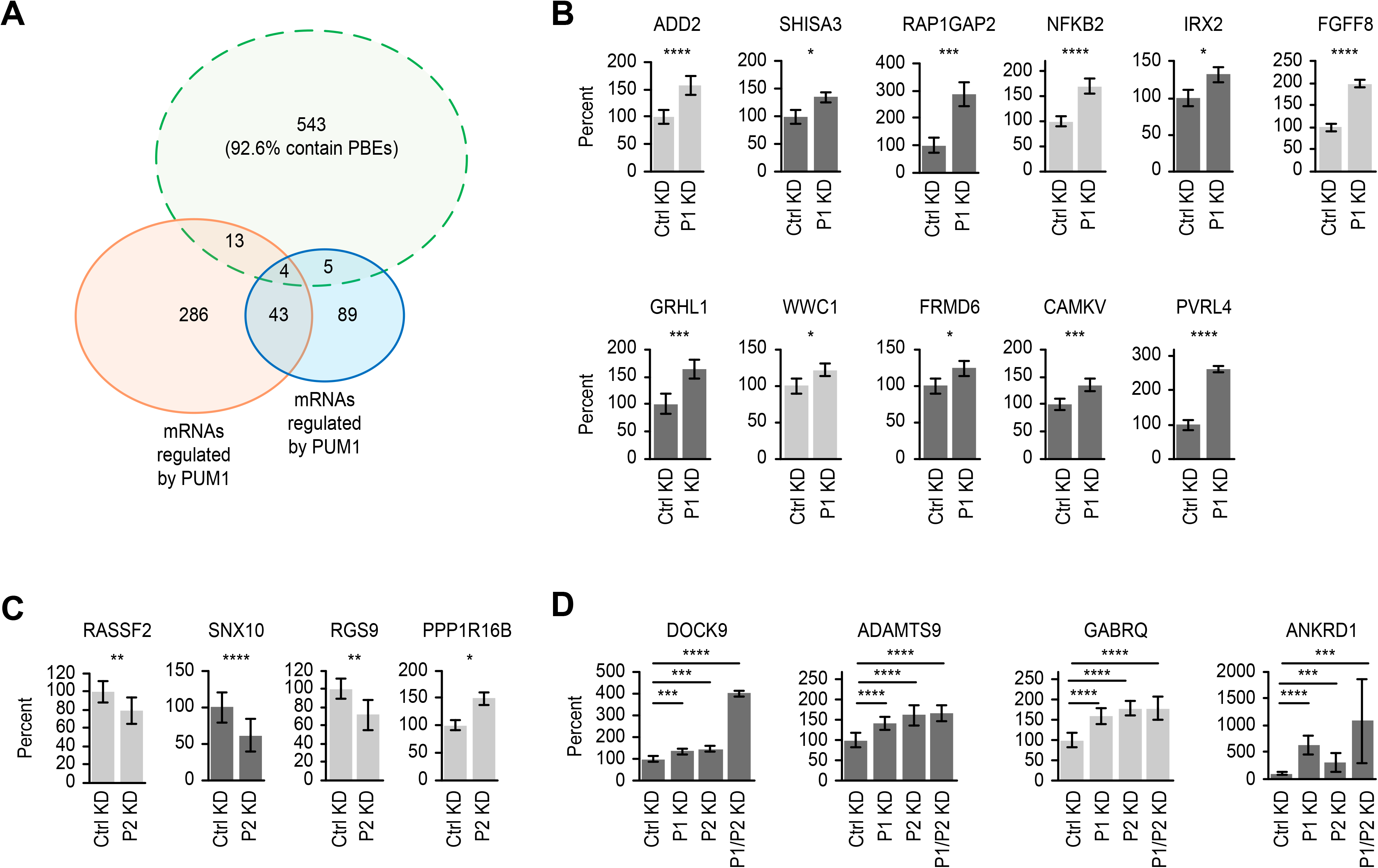
RT-qPCR validation of mRNAs regulated by PUM proteins that are enriched in TCam-2 cells compared to human somatic testis tissue. **A** Venn diagram showing 22 mRNAs identified as regulated by PUM1, PUM2 or both, selected from a pool of 565 mRNAs (92.6% of which was found to have PBE) found to be at least 6-fold enriched in TCam-2 cells, compared to somatic testis cells, as published by (29). **B** RT-qPCR validation of 11 out of 13 mRNAs selected as regulated by PUM1. **C** RT-qPCR validation of 4 out of 5 mRNAs selected as regulated by PUM2. **D** RT-qPCR validation of 4 out of 4 mRNAs selected as regulated by both PUM1 and PUM2. Dark grey histograms highlight mRNAs containing at least one classic PBE motif (UGUAHAUW), light grey histograms indicate mRNAs that do not contain any classic PBE. For all 3 repetitions of RT-qPCR, *ACTB* and *GAPDH* served as references. \**P*-value<0.05; \*\**P*-value<0.005; \*\*\**P*-value<0.0005; \*\*\*\**P*-value<0.00005.

To confirm that these 22 selected genes are under PUM1 and/or PUM2 regulation, we measured their expression by RT-qPCR in TCam-2 cells untreated or treated with *PUM1* siRNA, *PUM2* siRNA or both *PUM1* and *PUM2* siRNA. By this approach, we validated 19 of the 22 genes to be regulated by PUM proteins (**Fig. 6B, C, D** and **Table S8**) of which 11 (of 13) genes were regulated by PUM1 alone, 4 (of 5) mRNAs were regulated by PUM2 alone, and 4 (of 4) genes were regulated by both PUM1 and PUM2. The 11 genes regulated by PUM1 alone encode SHISA3, RAP1GAP2, NFKB2, IRX2, FGFR3, GRHL1, WWC1, FRMD6, CAMKV, PVRL4 and ADD2 (**Table S8**). Six of the eleven PUM1-regulated genes are associated with failure or cancer of the male as well as the female reproductive system. The 4 genes regulated by PUM2 alone encode RASSF2, SNX10, RGS9 and PPP1R16B (**Table S8**), which function in prostate tumour suppression, osteoporosis malignancy, nervous system development and endothelial cell proliferation, respectively. The 4 genes regulated by both PUM1 and PUM2 encode DOCK9, ADAMTS9, GABRQ and ANKRD1, which are involved in filopodia formation in cervical cancer, cell cycle regulation and ovary cancer progression, promotion of cell proliferation in hepatocellular carcinoma, downregulation of apoptosis, respectively. Three of these genes are associated with cancer of the reproductive system (**Table S8**).

We found that the majority of those PUM1- and PUM2-regulated targets are involved in cancer (16 among 19), including 10 in cancer of male or female reproductive system. PUM2-regulated PPP1R16B is functionally unique because it is the only PUM target that regulates phosphorylation. RT-qPCR validation of such a high proportion of genes indicates that our approach for PUM target identification was accurate.

Surprisingly, only 11 of the 19 validated genes contained at least one classic UGUAHAUW motif (**Fig. 6 B, C, D** dark grey bars), but all of them contain motifs with single nucleotide substitution in last 5 positions in comparison to classic PBE.

### Genes associated with male infertility in humans and/or mice are regulated by PUM proteins

We next sought to determine whether PUM-regulated genes in the male germ cell line, TCam-2, are associated with male infertility. To this end, from a list of 501 genes, which have been validated to cause infertility when mutated or disrupted in humans and/or mice (Matzuk & Lamb, 2008), we selected 11 genes that were also found in this study to be regulated by PUMs (**Fig. 7A** and **Table S8**). Upon *PUM1* and *PUM2* gene siRNA knockdown followed by RT-qPCR for these 11 genes, we validated 9 of them as PUM1/PUM2 targets. Of these 9 genes, 3 were regulated by PUM1 alone (**Fig. 7B**), 4 were regulated by PUM2 alone (**Fig. 7C**), and 2 were regulated by both PUM1 and PUM2 (**Fig. 7D**). We found, however, that only 2 of the PUM1-regulated *(FGFR2* and *NCOA6)* and 1 of the PUM1/PUM2-regulated *(LMTK2)* genes contained at least 1 classic UGUAHAUW motif (**Fig. 7B, C, D**), and all of them also contain motifs with single nucleotide substitution in last 5 positions in comparison to PBE. Dysfunction of all of the above 9 genes have been reported to be associated with male infertility or testis cancer (**Table S8**).

**Fig. 7.**
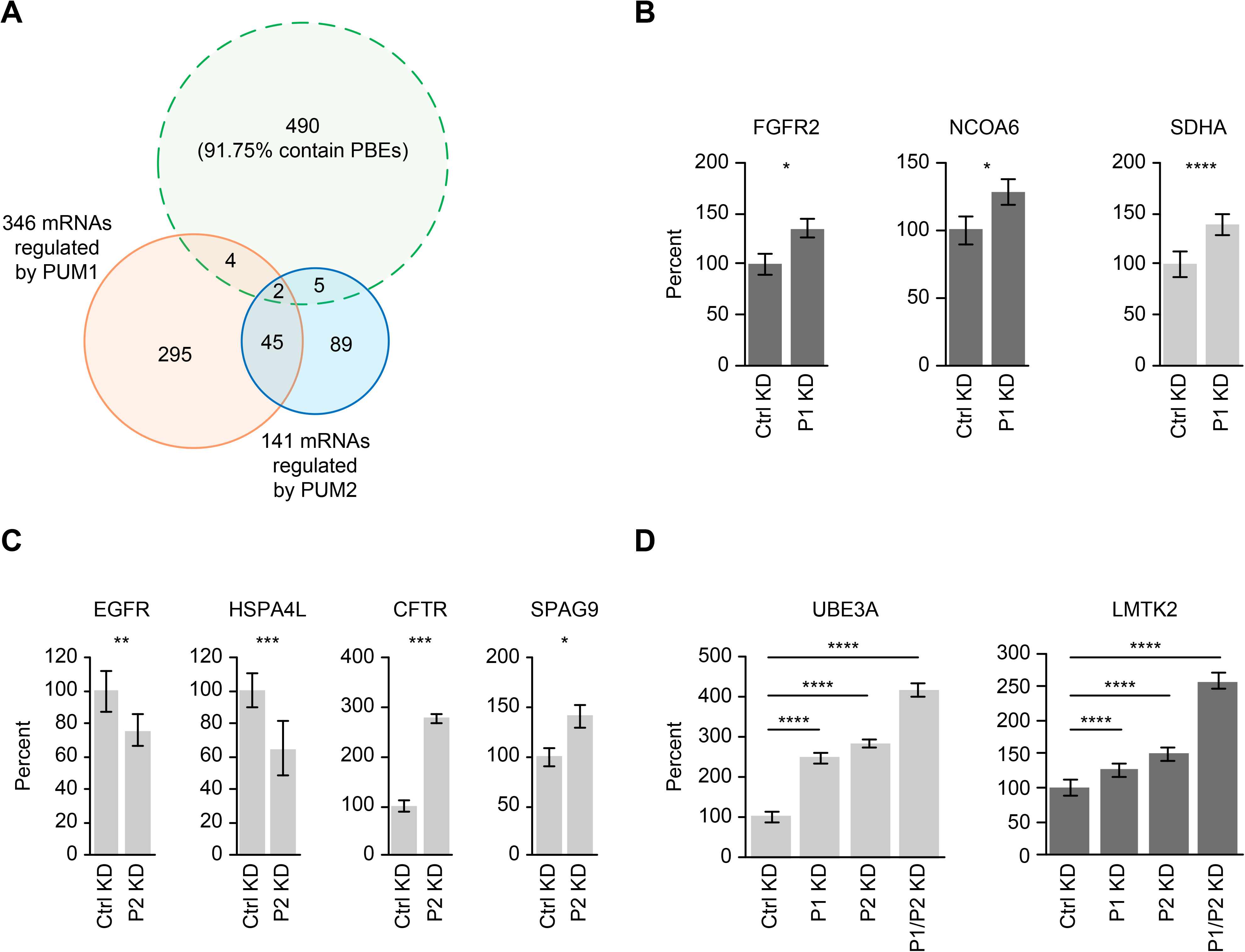
RT-qPCR validation of mRNAs regulated by PUM1 and/or PUM2, which are important for human and/or mouse male fertility. **A** Venn diagram showing 11 mRNAs identified as regulated by PUM1, PUM2 or both PUM1/PUM2 from a pool of 501 mRNAs (91.75% of which was found to have PBE) found to be involved in male infertility (Matzuk & Lamb, 2008). **B** RT-qPCR validation of 3 out of 4 mRNAs selected as regulated by PUM1. C RT-qPCR validation of 4 out of 5 mRNAs selected as regulated by PUM2. **D** RT-qPCR validation of 2 out of 2 mRNAs selected as regulated by PUM1 and PUM2. Dark grey histograms indicate mRNAs with at least one classic PBE motif (UGUAHAUW), light grey histograms indicate mRNAs that do not contain any classic PBE. For all 3 biological repetitions of RT-qPCR, *ACTB* and *GAPDH* served as references. \**P*-value<0.05; \*\**P*-value<0.005; \*\*\**P*-value<0.0005; \*\*\*\**P*-value<0.00005

## DISCUSSION

Considering high structural similarity of PUM1 and PUM2, it is still unresolved whether they form separate regulons in mammals. Here, for the first time, by combining RIP-Seq and siRNA-Seq data together with co-IP LC/MS identification of putative protein cofactors, RNA binding motif enrichment and GO analysis (for the first time each group of data originating from the same cells – TCam-2 cells) we obtained a model of regulatory networks distinct for PUM1 and PUM2. These networks are reminiscent of previously proposed regulons (Keene, 2007). Importantly, a global PUM-dependant gene expression regulation was not studied in germ cells, except *C. elegans* (Kershner & Kimble, 2010, Prasad et al., 2016).

It is important to note that several of our RIP-Seq identified targets overlapped with mRNAs previously identified in HeLa (Galgano et al., 2008) and HEK293 (Bohn et al., 2018, Hafner et al., 2010) cells, validating our results. However, it is also important to bear in mind that PUM-mediated activation or repression, or lack of PUM regulation may be cell-type-specific (Cottrell, Chaudhari et al., 2018). Therefore, we can expect only a partial target overlap when PUM targets from different types of cells are compared.

Furthermore, we found a much higher average representation of PBE-containing mRNAs that were selected as regulated by PUM1 and PUM2 based on combined analysis of RIP-Seq and PUM siRNA-Seq (96.80 and 99.80%, respectively), than in targets selected based on RIP-Seq alone (90.80 and 85.90%, respectively) or siRNA-Seq alone (59.68, 57.50%, respectively) which additionally validates our approach (**Fig. 2B**). We also analysed PBE motif distribution in the 5’UTR, CDS and 3’UTR of individual mRNA targets. We found that in mRNAs repressed by PUM1 or PUM2, PBEs were mostly localized in the 3’UTR, less frequently in CDS and rarely in the 5’UTR, as already reported (Bohn et al., 2018). Instead, in mRNAs positively regulated (activated/stabilized) by PUM1 or PUM2, PBE motifs were significantly more frequent in the 5’UTR (14.67% for PUM1 and 16.47% for PUM2) than in mRNAs negatively (repressed) by PUM1 and PUM2 (3.75 and 4.42%, respectively). However, this was not reported in studies on HEK293 cells (Bohn et al., 2018). Although this observation requires further studies, it may suggest that activation of these mRNAs by PUM proteins requires PBE localization in the vicinity of some 5’UTR translational signals. It is important to note that among PUM-regulated mRNAs, there are also a small number of targets with no PBE (approximately 3% PUM1- and below 1% PUM2-regulated). As mentioned above, PUM proteins may recognize motifs slightly different to the canonical UGUAHAUW (Jarmoskaite et al., 2019, Sajek et al., 2018). Such variant motifs were not evaluated in this study, and therefore, putative mRNA targets carrying such motifs were overlooked. It is also important to emphasize, that in our approach, PUM-regulated mRNAs whose level remained unchanged (do not undergo degradation or stabilization) were overlooked. PUM-regulated mRNA repression with no degradation but rather storage in P-bodies was recently suggested to be quite common in human HEK293 cells (Hubstenberger, Courel et al., 2017).

Interestingly, by using RIP-Seq approach we identified 30% of PUM1/PUM2-bound common targets. However, combination of RIP-Seq with siRNA knockdown to identify regulated targets, resulted in a decrease of common targets to 10%. We propose that this difference is due to the involvement of distinct regulatory factors for each PUM paralogue. It is worth emphasizing that we identified such regulatory factors – putative PUM-interacting protein cofactors which control different aspects of RNA metabolism (stability, localization, transport, splicing and expression regulation), whose interaction was RNA-mediated as well as protein cofactors whose interaction was RNA-independent. Substantial number of protein cofactors were PUM1- or PUM2-specific in both groups. The first group of RNA-dependent protein cofactors contains only RBPs, which was expected and validates our experiments as well as the analysis performed. However, RBPs were also significantly enriched in the second group representing RNA-independent protein-protein interactions. Such RBPs are likely to contain protein-protein interacting domains that bind PUM, as well as RNA-interacting domains that bind RNA. Finally, interactors with no RNA-binding domains might be important for the stabilization of ribonucleoprotein complexes, which are formed upon PUM protein binding specific mRNA targets.

Among the identified PUM putative protein interactors, we found five previously reported human PUM binding proteins, which validates our results. MATR3 and SEC16A, were previously identified in a high-throughput proteomic study in HeLa cells (Hein, Hubner et al., 2015). Another one is G3BP1, which is a stress granule assembly factor (Jain, Wheeler et al., 2016). The next one is the fragile X mental retardation protein (FMR1) and its autosomal homologous proteins, FXR1 and FXR2. FMR1 was previously shown to co-localize with PUM2 in rat neurons stress granules (Vessey, Vaccani et al., 2006). More recently, Zhang and co-workers reported that FMR1 interacts with PUM in the murine brain in an RNA-dependent manner (Zhang et al., 2017). In our study, FMR1 proteins were identified as both PUM RNA-dependent and independent interactors. Interactions with G3BP1 and FMR1 may suggest that PUM paralogues are components of stress granules not only in mammalian neurons (Vessey et al., 2006), but also in human germ cells. The presence of both PUM paralogues in stress granules suggests their involvement in RNA storage. Interestingly, PUM proteins were also found in P-bodies of HEK293 cells, which, according to a recent report (Hubstenberger et al., 2017), store high numbers of mRNAs.

Based on our results we propose that cooperation of such protein cofactors (mainly RBPs) with PUM1 or PUM2 enables regulation of selected groups of RNA targets which is responsible for a given metabolic pathway in TCam-2 cells. Notably, we found that a number of mRNAs which are enriched in TCam-2 cells compared to somatic gonadal tissue or cause infertility when mutated, are under the control of separate PUM1 or PUM2 regulons which is in line with their divergent functions. Additionally, each of them consists of sub-regulons. *NFKB2, FGFR3, FGFR2* and *NCOA6* are PUM1 targets (mRNAs are shown in **Table S7** functions and citations are in **Table S8**). *NFKB2* which is involved in aberrant activation of androgen receptor in prostate cancer cells, might be co-regulated by FXR2, NCL and HNRNPA1 proteins. FGFR3 which has a role in testis tumour development might be coregulated by a different set, SFPQ, FXR2 and HNRNPA1. FGFR2, which mutations were associated with hypospadias, might be co-regulated by SFPQ, NCL and HNRNPA1. Finally, NCOA6 which is involved in embryo implantation might be co-regulated by a large group of the following proteins: SFPQ, FXR1, FXR2, NCL, HNRNPA1, MATR3, PABPC1, IGFBP3, YBX1, NUDT21, IGF2BP1, PABPC4 and CPSF7. The same rule is observed in case of PUM2 targets involved in reproduction, *RASSF2, EGFR, CFTR* and *SPAG9* (**Table S7** and **S8**). RASSF2 which is a tumour-suppressor in prostate cancer mouse model might be coregulated by FXR1, FXR2, HNRNPA1, PTBP1, G3BP1, HNRNPF and SRSF7. EGFR, the signalling dysfunction of which was associated with human male infertility might be coregulated by FXR1, FXR2, NCL, HNRNPA1, G3BP2, G3BP1, FMR1, SRSF7 and SRSF1. CFTR, mutations of which are associated with male infertility, might be co-regulated by FXR1, NCL, HNRNPA1, MATR3, PABPC1, G3BP2, FMR1, SRSF7 and SRSF1. Finally, SPAG9, which stimulates prostate cancer cell proliferation might be co-regulated by FXR1, FXR2, NCL, HNRNPA1, MATR3, PABPC1, PTBP1, G3BP2, G3BP1, HNRNPF, FMR1 and SRSF7.

Therefore, we propose that identification of germ cell-associated groups of targets that are PUM1- or PUM2-specific indicates non-redundant roles of PUM paralogues in controlling processes of human reproduction. Notably, majority the PUM-regulated genes enriched in TCam-2 cells are genes involved in the development of several types of cancer, mostly of reproductive system (**Table S8**). This observation is in concordance with the fact that TCam-2 cells originate from seminoma testis germ cell tumour (de Jong et al., 2008).

Interestingly, although these PUM regulatory networks are distinct for each paralogue, they overlap at some points where PUM1 and PUM2 regulate some common targets and interact with some common protein cofactors (**Table S5** and **S6**). On the other hand, a PUM cofactor may regulate a specific pathway dependent on binding PUM1 or PUM2. For example, FXR2 may regulate endosome transport by binding PUM1 or Rho protein signal transduction by binding PUM2. Likewise, SFPQ may regulate cytosolic or endosome transport by binding PUM1 or endothelium development by binding PUM2.

The majority of selected PUM targets enriched in TCam-2 cells compared to gonadal tissue, have been reported to be involved in the regulation of the cell cycle, proliferation and apoptosis (**Table S8**), processes that are important for the maintenance of germ-line status and that are under precise regulation to ensure fertility. This functional profile is also in line with the recent suggestion that the evolutionarily original role of PUM proteins is regulation of stem cell self-renewal, including germline stem cells renewal (Bohn et al., 2018), which the above-mentioned three processes strongly influence.

Further studies of posttranscriptional mechanisms of gene expression regulation controlled by PUM proteins in the context of human germ cells are particularly important in light of the increasing problem of male and female infertility occurring worldwide in contemporary populations, as well as the increasing incidence of testis germ cell tumours in young men. In addition, it would be important to study the impact of PUM proteins on stem cell fate, growth and development, in the context of cancer and neurological disorders. This may provide insight into their diverse roles and enable future therapeutic strategies to target diseases arising from PUM and PUM-target dysfunctions.

## MATERIALS & METHODS

### RNA-immunoprecipitation with crosslinking and sequencing

For RIP analysis, TCam-2 cells were grown in 37°C and 5% CO_2_ in Roswell Park Memorial Institute (RPMI, Life Technologies 61870044) 1640 medium supplemented with 10% FBS (GE Healthcare HyClone SH30071) and 1% penicillin/streptomycin (Gibco 15140122). RIP-Seq experiments with UV cross-linking were performed using the Magna RIP^TM^ RNA-Binding Protein Immunoprecipitation Kit (17-700 Merck). Briefly, 100 μl of Magnetic A/G beads were coated with 12 μg anti-PUM1 (S-19, sc-65188 Santa Cruz Biotechnology), anti-PUM2 (K-14, sc-31535 Santa Cruz Biotechnology) antibody or IgG fraction from nonimmunized goat serum (G9759, Sigma Aldrich) for 45 min at room temperature (RT) in Magna RIP Wash Buffer. TCam-2 cells were washed twice with ice-cold PBS and subjected to UV cross-linking at 254 nm on a HEROLAB CL-1 Cross-linker for 30 s (0.015 J). For one RIP-Seq reaction, 2-3×10^6^ cells were lysed in 500 μl of Magna RIP Lysis Buffer for 30 min with rotation at 4°C. Lysates were centrifuged, and the supernatant was mixed with precoated beads suspended in washing buffer supplemented with protease and RNase inhibitors. The RIP reaction was held for 3 h at 4 °C on a rotator in a final volume of 1 ml. Then, magnetic beads were washed five times with Magna RIP Washing Buffer followed by treatment with proteinase K at 55 °C for 30 minutes. Total RNA was isolated from magnetic beads using a QIAGEN RNeasy Plus Micro Kit according to the manufacturer’s protocol, and RNA quality was checked on an Agilent Bioanalyzer using an RNA 6000 Nano Kit. RNA with a RIN value >7 was used for further steps. cDNA libraries for RNA-Seq analysis were prepared using Illumina TruSeq RNA Sample Prep V2, and subsequent next-generation sequencing was performed on an Illumina HiSeq 4000 platform by Macrogen INC. Sequencing was performed under the following conditions: Paired-End reads were 100 nt long, and >70 million reads/sample were obtained. RIP-Seq with anti-PUM1, PUM2, and IgG (negative control) were performed in triplicate. For TCam-2 transcriptome analysis, total RNA was isolated from 80% confluent 10 cm^2^ dishes using a QIAGEN RNeasy Plus Micro Kit. RNA quality control and RNA-Seq were performed as described above. An mRNA level that was at least 2-fold enriched (with adjusted *P*-value<0.05) in anti-PUM1/PUM2 co-IP, in comparison to the negative control (co-IP anti-IgG) and to the TCam-2 transcriptome level, was considered to be bound by PUM1 or PUM2.

### Western blot analysis

To check for PUM1 and PUM2 binding efficiency, SDS lysates from beads after co-IP were resolved on 8% SDS polyacrylamide gels and transferred to nitrocellulose membranes (BioRad). Membranes were blocked with 5% low-fat milk in TBS buffer supplemented with 0.1% Tween 20 (blocking buffer) at RT for 1 hour. Membranes were incubated with primary antibodies at 4°C overnight in blocking buffer. On the next day, membranes were washed 4 times in TBS buffer with 0.1% Tween 20 and incubated with horseradish peroxidase (HRP)-conjugated secondary antibodies at RT for 1 h in the same buffer. The following antibodies were used: goat anti-PUM1 (1:1000 Santa Cruz Biotechnology #S-19, sc-65188), goat anti-PUM2 (1:250 Santa Cruz Biotechnology #K-14, sc-31535), rabbit anti-actin beta (ACTB) (1:10000 Sigma Aldrich, A2066) and HRP-linked anti-goat (1:50000 Santa Cruz Biotechnology #sc-2020), as well as HRP-linked anti-rabbit (1:25000 Sigma Aldrich A0545). Next, membranes were washed twice in TBS buffer with 0.1% Tween 20, and then twice in TBS buffer. Clarity^TM^ ECL Western Blotting Substrate (BioRad) and the ChemiDoc Touch Imaging System (BioRad) were used for signal development and analysis. To check the silencing efficiency of PUM1 and PUM2, SDS lysates were prepared from cells 72 h posttransfection and analysed in the same way as lysates from beads.

### Bioinformatic analysis of PUM1- and PUM2-bound mRNAs

The Paired-End sequence reads obtained from the HiSeq 4000 platform were trimmed using the TRIMMOMATIC V0.35 tool with the following parameters: ILLUMINACLIP:TruSeq2:PE MINLEN:50, including quality filtration using SLIDINGWINDOW:10:25, MINLEN:50 parameter. Sequence reads that passed quality filters were mapped to the human reference genome (UCSC hg19) using TOPHAT(2.1.0) (Trapnell et al., 2009) with default parameters. Then, reads were counted using CUFFLINKS (2.2.1.0) followed by merging replicates with CUFFMERGE and calculating differential gene expression with CUFFDIFF (2.2.1.5).

For selection of mRNAs potentially bound to PUM1, PUM2 or both, the following criteria were used: 1) only mRNAs enriched in all 3 replicates; 2) at least 2-fold enrichment in PUM1 or PUM2 IPs, compared to IgG; and 3) at least 2-fold mRNA enrichment in comparison to TCam-2 transcriptome. To annotate mRNAs bound by PUM to their cell-specific functions and pathways, we performed GO analysis using BiNGO plug-in (version 3.0.3) (Maere et al., 2005) on Cytoscape platform (version 3.6.1) with functional annotation of biological process and molecular function, searched against TCam-2 cell line gene expression background derived from our RNA-Seq. Heatmaps were created using R (version 3.4.4) [R Core Team (2018). R: A language and environment for statistical computing. R Foundation for Statistical Computing, Vienna, Austria. URL https://www.R-project.org/.] and gplots R library. To identify PBEs in the 3’UTR of each PUM1- or PUM2-bound pool of mRNAs, we used the DREME motif discovery tool (ver.4.12.0), which enables the identification of short uninterrupted motifs that were enriched in our sequences compared with shuffled sequences. To search for PBE in whole mRNA PUM targets or their 5’UTR, CDS and 3’UTR, we used FIMO (ver.4.12.0), which enables scanning for individual matches for an input motif aligned to individual sequences. Whole mRNAs and their 3’UTR, 5’UTR and CDS sequences were downloaded from the RefSeq Genes Genomic Sequence database (Table browser: assembly: Feb. 2009(GRCh37/hg19); track: NCBI RefSeq).

### siRNA silencing of PUM proteins

TCam-2 cells were transfected with siRNA using PUM1 siRNA (sc-62912 Santa Cruz Biotechnology), PUM2 siRNA (sc-44773) containing 3 different siRNAs for each *PUM* gene (their sequences are in **Table S9**) or control siRNA-A (sc-37007) at the final 40 nM concentration using the NEON transfection/electroporation system (Thermo Fisher Scientific). Transfections were performed in Buffer R using 10 μl NEON tips. Subsequently, after transfection, cells were cultured in antibiotic-free RPMI 1640 (Gibco) supplemented with 10% FBS (HyClone) at 37 °C and 5% CO_2_ for 72 h. Transfection was performed in 3 biological replicates. RNA isolation was performed using a QIAGEN RNeasy Plus Micro Kit. RNA quality analysis was performed as described above, RNA with RIN>9 was used for cDNA library preparation, and subsequent sequencing was performed as described above. The knockdown efficiency of each replicate was analysed by western blot.

### Bioinformatic identification of mRNAs under regulation by PUM proteins

More than 80 million reads per sample obtained from the Illumina HiSeq 4000 platform were analysed as described above. If the mRNA level increased by at least 20% (adjusted *P*-value<0.05) under 70-90% *PUM* knockdown compared to negative siRNA control, the mRNA was considered to be under PUM repression. If the mRNA level decreased by at least 20%, it was considered to be significantly activated by PUM. We set the threshold at 20% as sufficient given that these changes were found in 3 biological replicates (adjusted *P*-value<0.05) and the protein silencing efficiency of PUM1 and PUM2 was high, (over 70% and 90%, respectively) (**Fig. S1C**).

### RT-qPCR analysis of mRNA regulation after PUM1 and PUM2 knockdown

To validate the targets regulated by PUM1 and PUM2, TCam-2 cells were transfected in 3 biological replicates with siRNA as described above. RNA from cells was isolated using TRIzol reagent (Gibco) according to the manufacturer’s protocol. ~1 μg of total RNA was treated with DNase I (D5307, Sigma-Aldrich) for 20 min at RT and reverse transcribed using the Maxima First-Strand cDNA Synthesis Kit (K1671, ThermoFisher Scientific) according to the manufacturer’s protocol. qPCR was performed on generated cDNA using Jump-Start Taq DNA Polymerase (D4184, Sigma-Aldrich), CFX96 Touch Real-Time PCR Detection System (BioRad) and SYBR Green dye (ThermoFisher Scientific) in 3 biological replicates with at least 5 technical replicates of each reaction. The list of primers used for RT-qPCR is shown in **Table S10**. All changes in mRNA levels upon PUM1 or PUM2 knockdown were normalized to ACTB and *GAPDH*.

### Mass spectrometry analysis after anti-PUM1 and anti-PUM2 immunoprecipitation

Six biological replicates of co-IPs (three performed without RNase A treatment, and another three with 100 mg/ml RNase A) with anti-PUM1, anti-PUM2 antibodies (including anti-IgG negative control to validate specificity of PUM1 and PUM2 antibodies and lack of cross reactivity) were performed as described above (**Table S1**). We used these antibodies in our MS/co-IP and RIP experiments. MS protein identification analysis was performed by MS Laboratory, IBB PAS, Warsaw. Briefly, proteins were directly digested on the beads and separated by liquid chromatography (LC) followed by MS measurement of peptides and their fragmentation spectra (LC-MS/MS) with a Q Exactive Hybrid Quadrupole-Orbitrap Mass Spectrometer (Thermo Scientific).

The bioinformatic protein identification analysis was performed as described: peak lists obtained from MS/MS spectra were identified using X! Tandem version X! Tandem Vengeance (2015.12.15.2), Andromeda version 1.5.3.4 and MS-GF+ version Beta (v10282). The search was conducted using SearchGUI version 3.2.23 (Barsnes & Vaudel, 2018).

Protein identification was conducted against a concatenated target/decoy (Elias & Gygi, 2010) version of the *Homo sapiens* OX=9606 (20316, 99.8%), cRAP (49, 0.2%), the complement of the UniProtKB (Apweiler, Bairoch et al., 2004) (version of [2017_06], 20365 (target) sequences). The decoy sequences were created by reversing the target sequences in SearchGUI. The identification settings were as follows: trypsin, specific, with a maximum of 1 missed cleavage of 30.0 ppm as MS1 and 0.1 Da as MS2 tolerances; fixed modifications: carbamidomethylation of C (+57.021464 Da), variable modifications: Oxidation of M (+15.994915 Da), fixed modifications during refinement procedure: carbamidomethylation of C (+57.021464 Da), variable modifications during refinement procedure: acetylation of protein N-term (+42.010565 Da), pyrrolidone from E (--18.010565 Da), pyrrolidone from Q (--17.026549 Da), pyrrolidone from carbamidomethylated C (--17.026549 Da). All specific algorithm settings are listed in the Certificate of Analysis available in the supplementary information.

Peptides and proteins were inferred from the spectrum identification results using PeptideShaker version 1.16.19 (Vaudel, Burkhart et al., 2015). Peptide spectrum matches (PSMs), peptides and proteins were validated at a 1.0% False Discovery Rate (FDR) estimated using the decoy hit distribution. All validation thresholds are listed in the Certificate of Analysis available in the supplementary information. List of identified proteins is shown in **Table S1** and **S6**. Proteins identified in 3 independent biological replicates of PUM1 or PUM2 IP and not identified in IgG IP were defined as PUM interactors (**Table S1** and **S6**).

### Bioinformatic construction of the PUM Regulon

Binding motifs of putative RNA-binding protein cofactors of PUM1 and PUM2 were obtained from RBPDB (http://rbpdb.ccbr.utoronto.ca/) CISP (http://cisbp-rna.ccbr.utoronto.ca/) and POSTAR2 (http://lulab.life.tsinghua.edu.cn/postar2/rbp2.php) databases (**Fig. 4D**). Motif enrichment analysis was performed on the identified mRNA targets of PUMs (**Fig. 1H**) by FIMO (http://meme-suite.org/doc/fimo.html) using a greater-than-average threshold (FIMO analysis with *P*-value <0.01; mRNAs for GO analysis bigger than average motif enrichment per sequence). mRNA groups regulated by PUM1 or PUM2 with the enrichment of the binding motif putative of RBP cofactors of the respective PUM (**Fig. 4**) were determined for each PUM-RBP co factor pair, in comparison to negative control mRNAs (not bound and not changed under PUM1 and PUM2 silencing). To avoid influence of sequence length, we selected negative sequences, which average length were similar (4177nt, in range 3000-16321) to PUM1 (4817nt, in range 449-16862) and PUM2 (5442nt, in range 412-16862) regulated mRNAs. As the next step, GO analysis of biological processes on identified groups was performed using ClueGO version 2.5.2 (Bindea, Mlecnik et al., 2009). Pathways with *P*-values≤ 0.05 were considered significant. Visualization of the regulon was performed using Cytoscape platform version 3.6.1.

## DATA AVAILABILITY

Processed and raw data for RIP-Seq and RNA-Seq experiments described here are available from the Gene Expression Omnibus (accession GSE123016). The mass spectrometry proteomics data have been deposited to the ProteomeXchange Consortium via the PRIDE (Vizcaino, Csordas et al., 2016) partner repository with the dataset identifier PXD011948 and 10.6019/PXD011948. During the review process, the data can be accessed with the following credentials upon login to the PRIDE website (http://www.ebi.ac.uk/pride/archive/login): Username, Password: [OigS3Vy7].

## ACKNOWLEDGMENTS

We thank members of the JJ lab for reading the manuscript.

## AUTHOR CONTRIBUTIONS

Conceptualization, M.J.S., E.I., M.S. and J.J.; methodology, M.J.S., E.I., M.S., A.S., D.M.J., K.K-Z., T.W., L.H.; investigation, M.J.S, E.I., M.S., K.K-Z.; formal analysis, M.J.S., E.I., M.S., T.W., A.S.; writing – original Draft, M.J.S., M.S. and J.J.; writing – review & editing, M.J.S., M.S., E.I. and J.J.; funding acquisition, M.J.S., J.J. and K.K-Z; resources, M.F. and J.J.; supervision, J.J.

---

**Figure S1 Related to Figure 1.**
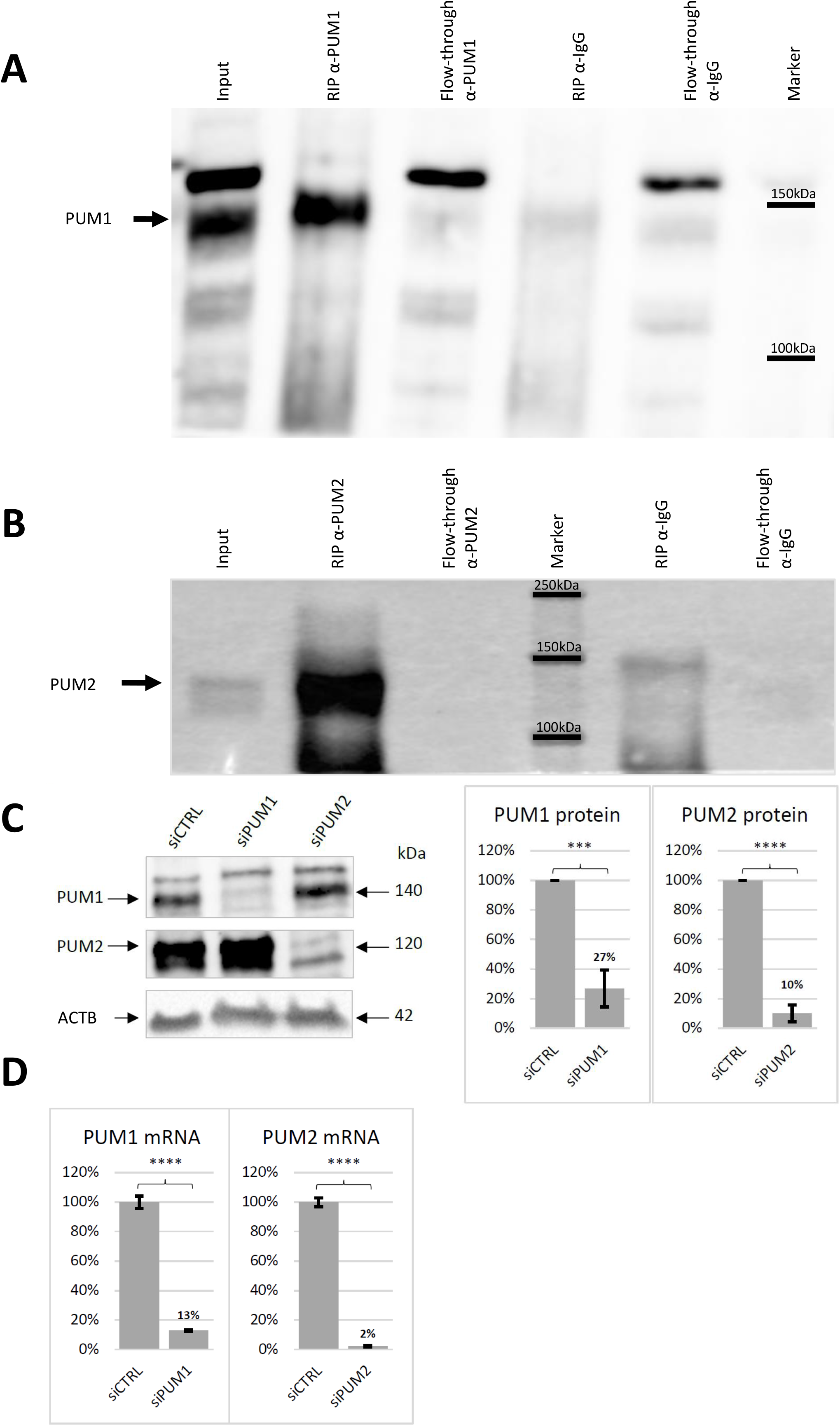
Western blot (WB) and RT-qPCR analyses for the estimation of PUM binding efficiency to beads and PUM knockdown efficiency for RNA-Seq. **A** WB detection of PUM1 on beads after co-immunoprecipitation (co-IP) using anti-PUM1 antibody. **B** WB detection of PUM2 on beads after Co-IP using anti-PUM2 antibody. **C** Representative WB for PUM1 and PUM2 after siRNA knockdown (left panel) and histograms showing quantitation of protein knockdown from 3 biological repetitions. For quantitative analyses, ACTB was treated as a reference. **D** Histograms showing quantitation of PUM1 and PUM2 mRNA knockdown from 3 biological repetitions, in which ACTB and GAPDH mRNAs were treated as references. *** Pvalue< 0.0005; **** P-value<0.00005.

**Figure S2 Related to Figure 1 and 6.**
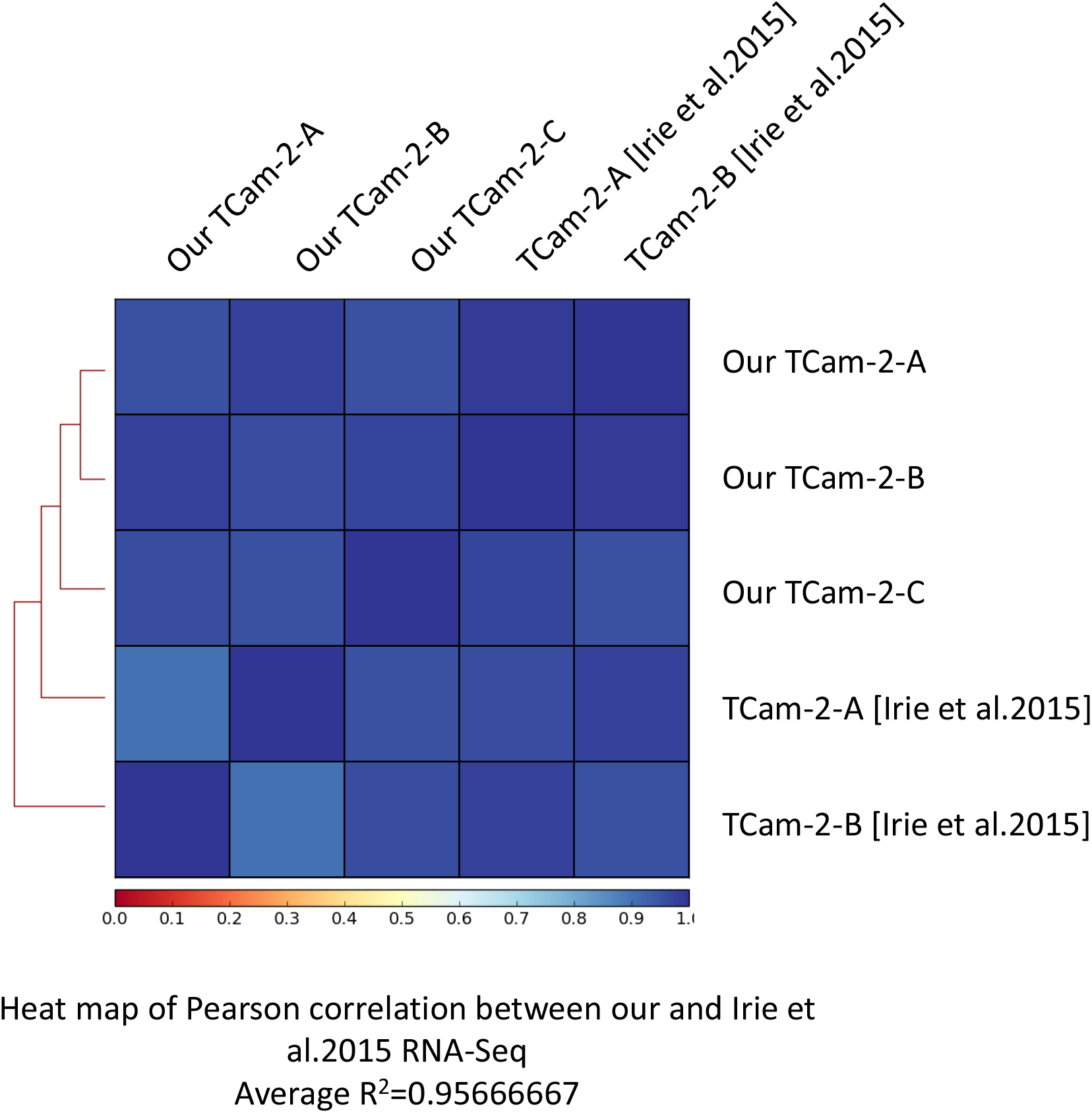
Heat map of Pearson correlation for comparison of the TCam-2 transcriptome obtained in this study with the TCam-2 transcriptome published by (Irie et al., 2015).

**Figure S3 Related to Figure 1.**
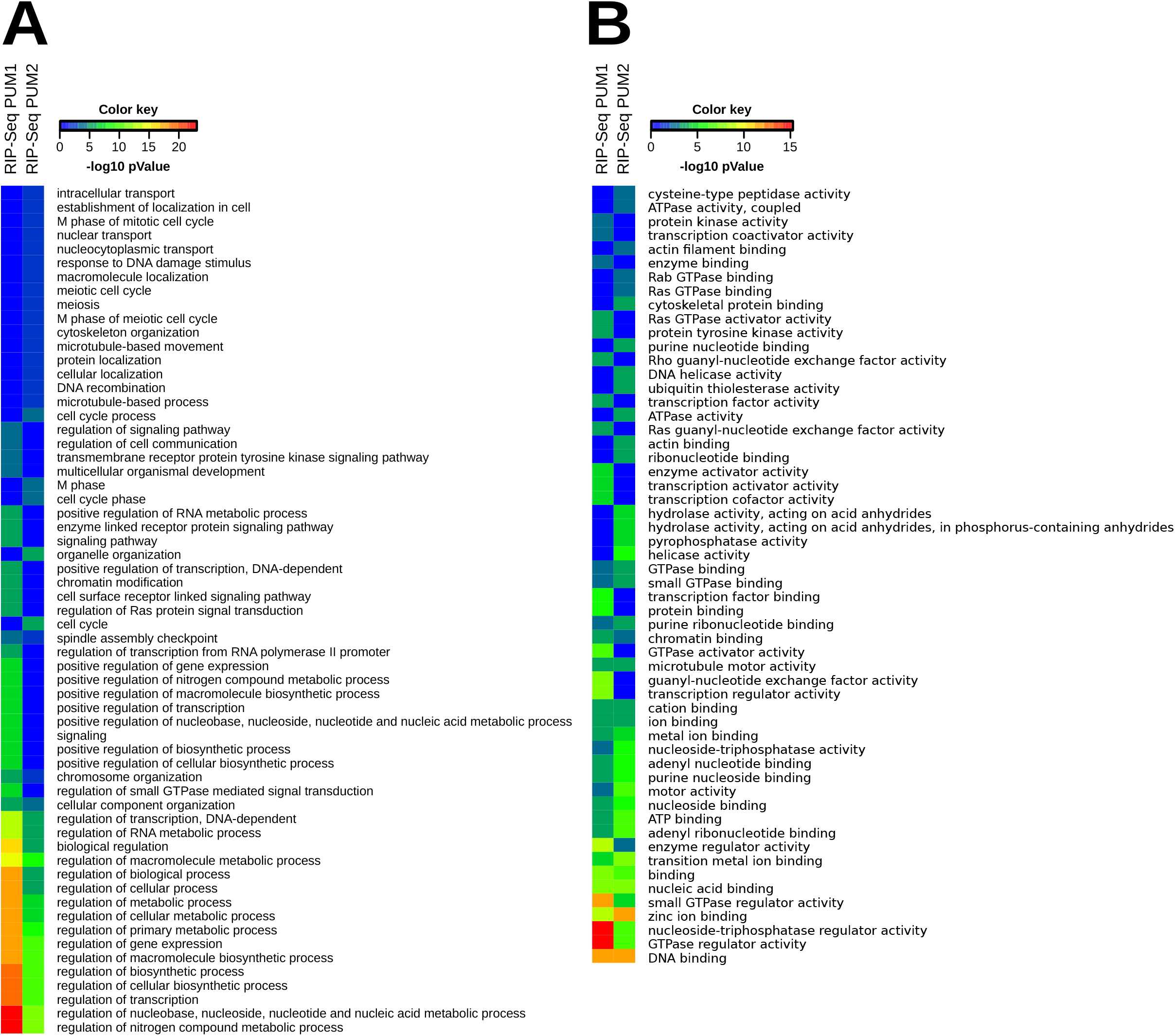
Gene Ontology analysis of mRNAs bound to PUM1 and PUM2 as identified by the RIP-Seq approach. **A** Heatmap of BiNGO analysis of TOP40 Biological Processes of mRNAs selected as bound to PUM1 and PUM2. **B** Heatmap of BiNGO analysis of TOP40 Molecular Functionsof mRNAs selected as bound to PUM1 and PUM2. The detailed results of the GOanalysis are presented in **Table S3**.

**Figure S4 Related to Figure 1.**
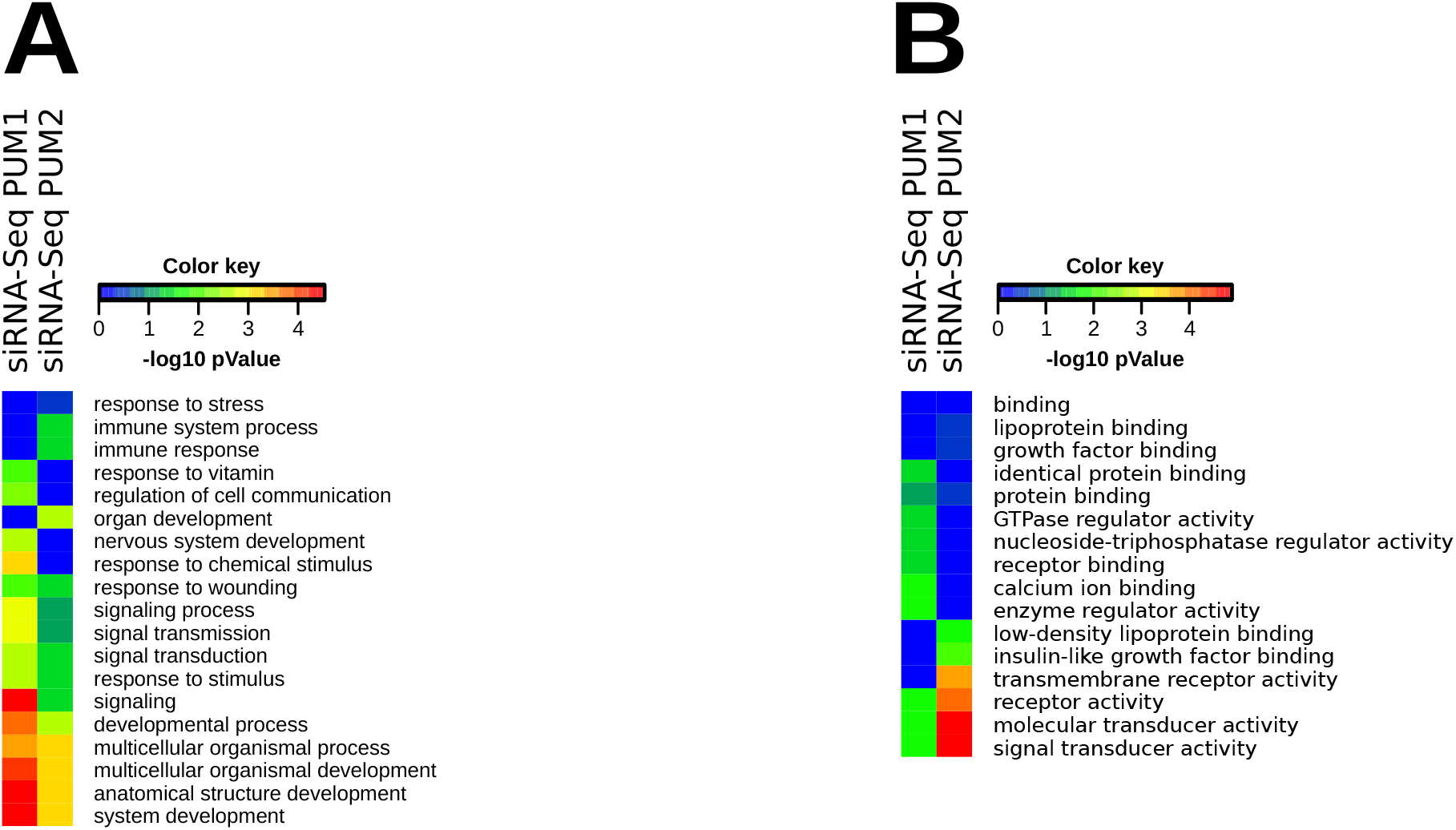
Gene Ontology analysis of mRNAs up-and downregulated upon PUM1 and/or PUM2 siRNA knockdown (PUM1 or PUM2 siRNA-Seq) **A** Heatmap of BiNGO analysis of TOP15 Biological Processesof mRNAs selected as up-anddownregulated upon PUM1 or PUM2 siRNA knockdown. **B** Heatmap of BiNGO analysis of TOP10 Molecular Functionsof mRNAs selected as up-anddownregulated upon PUM1 or PUM2 siRNA. The detailed results of the GO analysis are presented in **Table S3**.

**Figure S5 Related to Figure 1.**
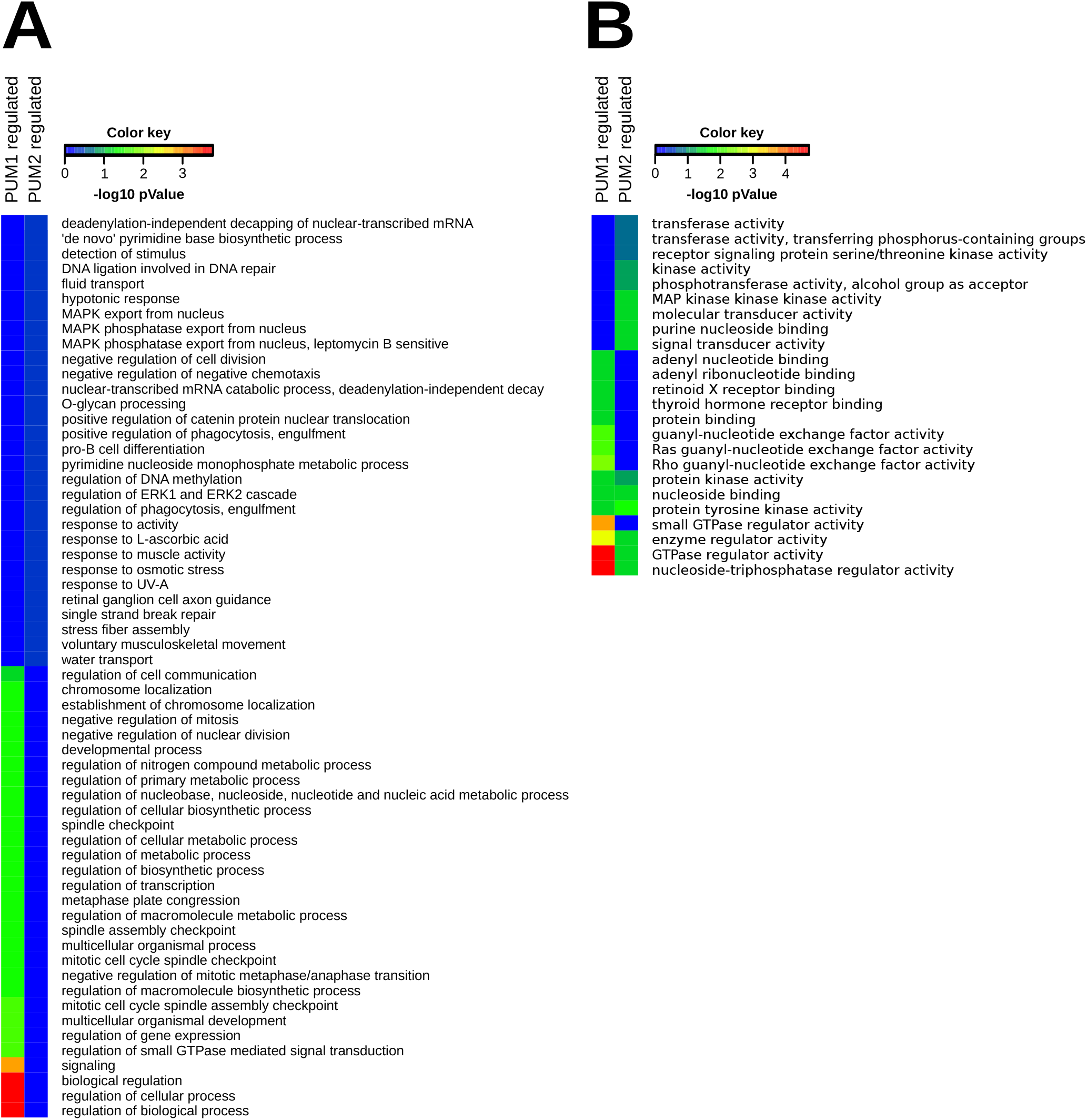
Gene Ontology analysis of PUMl-and PUM2-regulated targets. **A** Heatmap of BiNGO analysis of TOP30 Biological Processesof mRNAs regulated by PUM1 and PUM2. **B** Heatmap ofBiNGO analysis of TOP15 Molecular Functions of mRNAs regulated by PUM1 and PUM2. The detailed results of the GOanalysis are presented in **Table S3**.

## REFERENCES

Apweiler R, Bairoch A, Wu CH, Barker WC, Boeckmann B, Ferro S, Gasteiger E, Huang H, Lopez R, Magrane M, Martin MJ, Natale DA, O’Donovan C, Redaschi N, Yeh LS (2004) UniProt: the Universal Protein knowledgebase. Nucleic Acids Res 32: D115–9

Barsnes H, Vaudel M (2018) SearchGUI: A Highly Adaptable Common Interface for Proteomics Search and de Novo Engines. J Proteome Res 17: 2552–2555

Bindea G, Mlecnik B, Hackl H, Charoentong P, Tosolini M, Kirilovsky A, Fridman WH, Pages F, Trajanoski Z, Galon J (2009) ClueGO: a Cytoscape plug-in to decipher functionally grouped gene ontology and pathway annotation networks. Bioinformatics 25: 1091–3

Bohn JA, Van Etten JL, Schagat TL, Bowman BM, McEachin RC, Freddolino PL, Goldstrohm AC (2018) Identification of diverse target RNAs that are functionally regulated by human Pumilio proteins. Nucleic Acids Res 46: 362–386

Chen D, Zheng W, Lin A, Uyhazi K, Zhao H, Lin H (2012) Pumilio 1 suppresses multiple activators of p53 to safeguard spermatogenesis. Curr Biol 22: 420–5

Cottrell KA, Chaudhari HG, Cohen BA, Djuranovic S (2018) PTRE-seq reveals mechanism and interactions of RNA binding proteins and miRNAs. Nat Commun 9: 301

de Jong J, Stoop H, Gillis AJ, Hersmus R, van Gurp RJ, van de Geijn GJ, van Drunen E, Beverloo HB, Schneider DT, Sherlock JK, Baeten J, Kitazawa S, van Zoelen EJ, van Roozendaal K, Oosterhuis JW, Looijenga LH (2008) Further characterization of the first seminoma cell line TCam-2. Genes Chromosomes Cancer 47: 185–96

Elias JE, Gygi SP (2010) Target-decoy search strategy for mass spectrometry-based proteomics. Methods Mol Biol 604: 55–71

Fredericks AM, Cygan KJ, Brown BA, Fairbrother WG (2015) RNA-Binding Proteins: Splicing Factors and Disease. Biomolecules 5: 893–909

Galgano A, Forrer M, Jaskiewicz L, Kanitz A, Zavolan M, Gerber AP (2008) Comparative analysis of mRNA targets for human PUF-family proteins suggests extensive interaction with the miRNA regulatory system. PLoS One 3: e3164

Gerber AP, Herschlag D, Brown PO (2004) Extensive association of functionally and cytotopically related mRNAs with Puf family RNA-binding proteins in yeast. PLoS Biol 2: E79

Gerstberger S, Hafner M, Tuschl T (2014) A census of human RNA-binding proteins. Nat Rev Genet 15: 829–45

Goldstrohm AC, Hall TMT, McKenney KM (2018) Post-transcriptional Regulatory Functions of Mammalian Pumilio Proteins. Trends Genet

Hafner M, Landthaler M, Burger L, Khorshid M, Hausser J, Berninger P, Rothballer A, Ascano M, Jr., Jungkamp AC, Munschauer M, Ulrich A, Wardle GS, Dewell S, Zavolan M, Tuschl T (2010) Transcriptome-wide identification of RNA-binding protein and microRNA target sites by PAR-CLIP. Cell 141: 129–41

Hein MY, Hubner NC, Poser I, Cox J, Nagaraj N, Toyoda Y, Gak IA, Weisswange I, Mansfeld J, Buchholz F, Hyman AA, Mann M (2015) A human interactome in three quantitative dimensions organized by stoichiometries and abundances. Cell 163: 712–23

Hubstenberger A, Courel M, Benard M, Souquere S, Ernoult-Lange M, Chouaib R, Yi Z, Morlot JB, Munier A, Fradet M, Daunesse M, Bertrand E, Pierron G, Mozziconacci J, Kress M, Weil D (2017) P-Body Purification Reveals the Condensation of Repressed mRNA Regulons. Mol Cell 68: 144–157 e5

Ibtisham F, Wu J, Xiao M, An L, Banker Z, Nawab A, Zhao Y, Li G (2017) Progress and future prospect of in vitro spermatogenesis. Oncotarget 8: 66709–66727

Irie N, Weinberger L, Tang WW, Kobayashi T, Viukov S, Manor YS, Dietmann S, Hanna JH, Surani MA (2015) SOX17 is a critical specifier of human primordial germ cell fate. Cell 160: 253–68

Jain S, Wheeler JR, Walters RW, Agrawal A, Barsic A, Parker R (2016) ATPase-Modulated Stress Granules Contain a Diverse Proteome and Substructure. Cell 164: 487–98

Jarmoskaite I, Denny SK, Vaidyanathan PP, Becker WR, Andreasson JOL, Layton CJ, Kappel K, Shivashankar V, Sreenivasan R, Das R, Greenleaf WJ, Herschlag D (2019) A Quantitative and Predictive Model for RNA Binding by Human Pumilio Proteins. Mol Cell 74: 966–981 e18

Jaruzelska J, Kotecki M, Kusz K, Spik A, Firpo M, Reijo Pera RA (2003) Conservation of a Pumilio-Nanos complex from Drosophila germ plasm to human germ cells. Dev Genes Evol 213: 120–6

Keene JD (2007) RNA regulons: coordination of post-transcriptional events. Nat Rev Genet 8: 533–43

Kershner AM, Kimble J (2010) Genome-wide analysis of mRNA targets for Caenorhabditis elegans FBF, a conserved stem cell regulator. Proc Natl Acad Sci U S A 107: 3936–41

Kusz-Zamelczyk K, Sajek M, Spik A, Glazar R, Jedrzejczak P, Latos-Bielenska A, Kotecki M, Pawelczyk L, Jaruzelska J (2013) Mutations of NANOS1, a human homologue of the Drosophila morphogen, are associated with a lack of germ cells in testes or severe oligo-astheno-teratozoospermia. J Med Genet 50: 187–93

Maere S, Heymans K, Kuiper M (2005) BiNGO: a Cytoscape plugin to assess overrepresentation of gene ontology categories in biological networks. Bioinformatics 21: 3448–9

Mak W, Fang C, Holden T, Dratver MB, Lin H (2016) An Important Role of Pumilio 1 in Regulating the Development of the Mammalian Female Germline. Biol Reprod 94: 134

Matzuk MM, Lamb DJ (2008) The biology of infertility: research advances and clinical challenges. Nat Med 14: 1197–213

Moore FL, Jaruzelska J, Fox MS, Urano J, Firpo MT, Turek PJ, Dorfman DM, Pera RA (2003) Human Pumilio-2 is expressed in embryonic stem cells and germ cells and interacts with DAZ (Deleted in AZoospermia) and DAZ-like proteins. Proc Natl Acad Sci U S A 100: 538–43

Morris AR, Mukherjee N, Keene JD (2008) Ribonomic analysis of human Pum1 reveals cis-trans conservation across species despite evolution of diverse mRNA target sets. Mol Cell Biol 28: 4093–103

Mukherjee N, Wessels HH, Lebedeva S, Sajek M, Ghanbari M, Garzia A, Munteanu A, Yusuf D, Farazi T, Hoell JI, Akat KM, Akalin A, Tuschl T, Ohler U (2019) Deciphering human ribonucleoprotein regulatory networks. Nucleic Acids Res 47: 570–581

O’Flynn O’Brien KL, Varghese AC, Agarwal A (2010) The genetic causes of male factor infertility: a review. Fertil Steril 93: 1–12

Prasad A, Porter DF, Kroll-Conner PL, Mohanty I, Ryan AR, Crittenden SL, Wickens M, Kimble J (2016) The PUF binding landscape in metazoan germ cells. RNA 22: 1026–43

Ray D, Kazan H, Cook KB, Weirauch MT, Najafabadi HS, Li X, Gueroussov S, Albu M, Zheng H, Yang A, Na H, Irimia M, Matzat LH, Dale RK, Smith SA, Yarosh CA, Kelly SM, Nabet B, Mecenas D, Li W et al. (2013) A compendium of RNA-binding motifs for decoding gene regulation. Nature 499: 172–7

Reijo R, Lee TY, Salo P, Alagappan R, Brown LG, Rosenberg M, Rozen S, Jaffe T, Straus D, Hovatta O, et al. (1995) Diverse spermatogenic defects in humans caused by Y chromosome deletions encompassing a novel RNA-binding protein gene. Nat Genet 10: 383–93

Sajek M, Janecki DM, Smialek MJ, Ginter-Matuszewska B, Spik A, Oczkowski S, Ilaslan E, Kusz-Zamelczyk K, Kotecki M, Blazewicz J, Jaruzelska J (2018) PUM1 and PUM2 exhibit different modes of regulation for SIAH1 that involve cooperativity with NANOS paralogues. Cell Mol Life Sci

Santos MG, Machado AZ, Martins CN, Domenice S, Costa EM, Nishi MY, Ferraz-de-Souza B, Jorge SA, Pereira CA, Soardi FC, de Mello MP, Maciel-Guerra AT, Guerra-Junior G, Mendonca BB (2014) Homozygous inactivating mutation in NANOS3 in two sisters with primary ovarian insufficiency. Biomed Res Int 2014: 787465

Spassov DS, Jurecic R (2003) The PUF family of RNA-binding proteins: does evolutionarily conserved structure equal conserved function? IUBMB Life 55: 359–66

Sternburg EL, Estep JA, Nguyen DK, Li Y, Karginov FV (2018) Antagonistic and cooperative AGO2-PUM interactions in regulating mRNAs. Sci Rep 8: 15316

van de Geijn GJ, Hersmus R, Looijenga LH (2009) Recent developments in testicular germ cell tumor research. Birth Defects Res C Embryo Today 87: 96–113

Vaudel M, Burkhart JM, Zahedi RP, Oveland E, Berven FS, Sickmann A, Martens L, Barsnes H (2015) PeptideShaker enables reanalysis of MS-derived proteomics data sets. Nat Biotechnol 33: 22–4

Vessey JP, Vaccani A, Xie Y, Dahm R, Karra D, Kiebler MA, Macchi P (2006) Dendritic localization of the translational repressor Pumilio 2 and its contribution to dendritic stress granules. J Neurosci 26: 6496–508

Vizcaino JA, Csordas A, Del-Toro N, Dianes JA, Griss J, Lavidas I, Mayer G, Perez-Riverol Y, Reisinger F, Ternent T, Xu QW, Wang R, Hermjakob H (2016) 2016 update of the PRIDE database and its related tools. Nucleic Acids Res 44: 11033

Wang X, McLachlan J, Zamore PD, Hall TM (2002) Modular recognition of RNA by a human pumilio-homology domain. Cell 110: 501–12

Weidmann CA, Goldstrohm AC (2012) Drosophila Pumilio protein contains multiple autonomous repression domains that regulate mRNAs independently of Nanos and brain tumor. Mol Cell Biol 32: 527–40

Xu EY, Chang R, Salmon NA, Reijo Pera RA (2007) A gene trap mutation of a murine homolog of the Drosophila stem cell factor Pumilio results in smaller testes but does not affect litter size or fertility. Mol Reprod Dev 74: 912–21

Zhang M, Chen D, Xia J, Han W, Cui X, Neuenkirchen N, Hermes G, Sestan N, Lin H (2017) Post-transcriptional regulation of mouse neurogenesis by Pumilio proteins. Genes Dev 31: 1354–1369

